# Cultivation of widespread *Bathyarchaeia* reveals a novel methyltransferase system utilizing lignin-derived aromatics

**DOI:** 10.1101/2022.10.23.513425

**Authors:** Tiantian Yu, Haining Hu, Xianhong Zeng, Yinzhao Wang, Donald Pan, Fengping Wang

**Affiliations:** School of Oceanography, Shanghai Jiao Tong University, 200240 Shanghai, China; State Key Laboratory of Microbial Metabolism, School of Life Sciences and Biotechnology, Shanghai Jiao Tong University, 200240 Shanghai, China; Southern Marine Science and Engineering Guangdong Laboratory (Zhuhai), Zhuhai, Guangdong, China

## Abstract

Anaerobic lignin degradation is a major process in the global carbon cycle that would significantly influence estimates of carbon flux in both terrestrial and marine ecosystems. The ubiquitous *Bathyarchaeia*, one of the most abundant taxa in marine sediments, have been proposed to be key players in this process. However, the mechanism of Bathyarchaeial lignin degradation is unclear due to the lack of cultured strains. Here we report the cultivation of Candidatus *Marisediminiarchaeum ligniniphilus* DL1YTT001, a Bathyarchaeial representative from nearshore marine sediments that can grow with lignin as the sole organic carbon source under mesophilic and anaerobic conditions. Strain DL1YTT001 possesses and highly expresses a novel and specific methyltransferase system for O-demethylation of lignin-derived methoxylated aromatic compounds (ArOCH_3_). The key gene, methyltransferase 1 (MtgB), is not homologous to any other lineages. Enzymatic activity was confirmed through the heterologous expression of the MtgB gene, showing O-demethylation activity with guaiacol as the substrate. Considering that Bathyarchaeial lineages carrying this specific methyltransferase system are widely distributed in diverse anoxic environments, especially lignin-rich nearshore sediments, *Bathyarchaeia*-mediated O-demethylation is likely a key step in global anaerobic lignin remineralization.

## Background

Marine sediments are one of largest organic carbon reservoirs on Earth with nearshore sediments contributing up to 45% of total organic carbon buried within marine sediments^1^. Biomineralization of sedimentary organic matter is performed by microbial respiration with oxygen as the primary electron acceptor^2^. In nearshore sediments with high organic matter loading, oxygen is quickly exhausted within the top a few millimeters to centimeters^3, 4^, leading to an extensive anaerobic zone where a large proportion of organic carbon persists, including a considerable amount of recalcitrant lignin^5–7^. Lignin, a complex aromatic polymer highly resistant to biological degradation, composes approximately 25% of the dry weight of vascular plants which comprise a large proportion (20-50%) of the organic carbon deposited into nearshore sediments^8,9^

Methoxylated aromatic compounds (ArOCH_3_) are a major component of lignin with ~3% of the total carbon content of lignin being in the form of methoxyl groups^10, 11^. Under anaerobic conditions, acetogenic Bacteria can demethylate from ArOCH_3_ and utilize the methyl groups as a source of energy^12^. The demethylation starts with cleavage of the ether bond linking the phenyl and methyl group, follow by transfer of the methyl group to an acceptor tetrahydrofolate (H_4_F). Methyltransferases have been characterized in acetogenic Bacteria such as *Moorella thermoacetica*^13^, *Acetobacterium dehalogenans*^14^, *Sporomusa termitida*^15^, and *Acetobacterium woodii*^12^. In recent years, these Bacterial-type methyltransferase systems were also detected in Archaea, including an acetogenic archaeon *Archaeoglobus fulgidus* and a methanogenic archaeon *Methermicoccus shengliensis*^16,17^ Similar genes were also found in the genomes of some uncultured Archaea such as *Bathyarchaeia, Lokiarchaeia, Methanomethylicia, Korarchaeia, Helarchaeota* and *Nezhaarchaeales*^17^.

The uncultured *Bathyarchaeia* are estimated to be among the most abundant microorganisms on Earth^18, 19^ They are ubiquitously distributed in nearly all types of anaerobic environments, particularly in anaerobic nearshore sediments^19–21^. The members of *Bathyarchaeia* show high diversity and were previously classified into 25 subgroups^21^, now assigned into seven orders based on the Genome Taxonomy Database (GTDB)^22, 23^. Moreover, its members have the genomic potential for diverse carbon metabolisms, such as carbon fixation, acetogenesis, alkane metabolism, and fermentation of a wide variety of organic substrates^18, 19, 21, 22, 24, 25^, suggesting that they may play an important role in global carbon cycling. We prior study found that the addition of kraft lignin to nearshore sediment samples significantly stimulated the growth of *Bathyarchaeia*^26^. During the enrichment with lignin, assimilation of inorganic carbon into Archaeal lipids was observed, suggesting an organoautotrophic lifestyle of *Bathyarchaeia*. This also implies that the *Bathyarchaeia* play an important role in lignin degradation in anoxic nearshore sediments. However, no known genes encoding lignin polymer depolymerization enzymes have been found in metagenomic assembled genomes (MAGs) of *Bathyarchaeia*. We hypothesized that the mode of lignin metabolism of *Bathyarchaeia* may be similar to that described for acetogenic Bacteria with the capability of O-demethylation of ArOCH_3_^26^. In this study, we successfully cultured an archaeon of *Bathyarchaeia* on lignin-containing media. Based on genomic, transcriptomic, proteomic, and enzymological evidence, we propose a novel and *Bathyarchaeia*-specific methyltransferase system for O-demethylation from ArOCH_3_.

## Result

### Cultivation and characterization of a *Bathyarchaeia* archaeon

In 2018, we enriched *Bathyarchaeia* from nearshore sediment samples by adding lignin as the sole organic carbon source, followed by one year of incubation^26^. The enrichment culture was transferred to artificial seawater medium supplemented with 0.5% kraft lignin and 50 mM sodium bicarbonate and incubated at 35°C. The growth of *Bathyarchaeia* was detected by quantitative PCR after a two month lag phase. Since then, Bathyarchaeial growth has been maintained in our lab for more than five years by consecutive 1:10 transfers at 2-3 months intervals. One strain of *Bathyarchaeia* named DL1YTT001 was dominant in the enrichment culture and comprised approximately 60% of the total abundance according to analysis of 16S rRNA genes amplicon sequencing (2021/12, Fig. 1a and Table S1). This DL1YTT001 enrichment culture also contains some Bacterial groups, including *Desulfobacteraceae (Desulfobacterota*) 13.22%, *Spirochaetaceae (Spirochaetota*) 10.50%, *Desulfovibrionaceae (Desulfobacterota*)5.46%, *Deferribacteraceae (Deferribacterota*) 2.18%, *Synergistaceae (Synergistota*)2.05%, *Christensenellaceae (Firmicutes*) 1.64%*etc*. The 16S rRNA gene copy numbers of the strain DL1YTT001 increased 37-fold in 20 days from ~2.7×10^7^ to 1.7×10^8^ gene copies/ml, and its doubling time was estimated to be approximately 3.8 days (Fig. 1b).

**Fig. 1.**
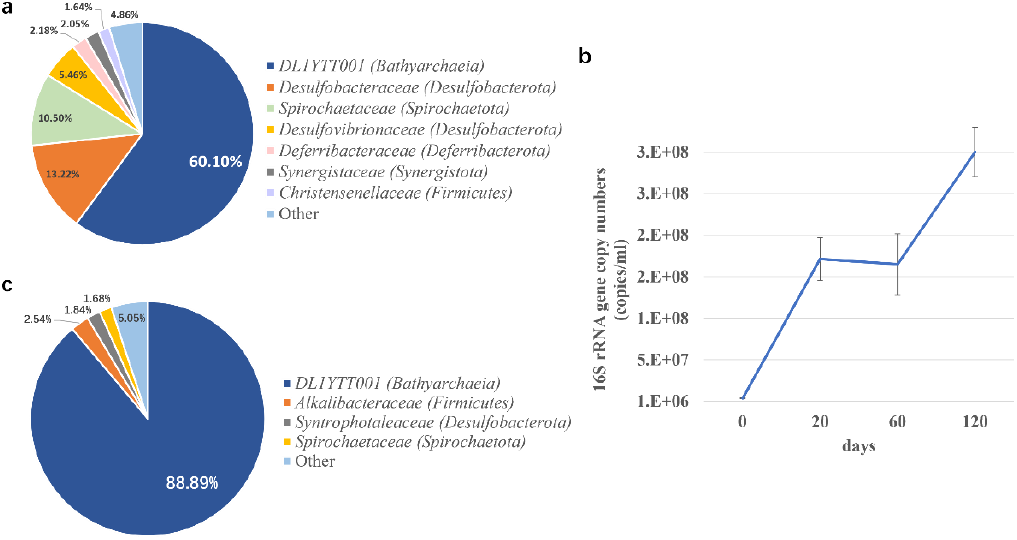
Fluorescence images and growth curves of the cultured Bathyarchaeotial strain DL1YTT001 and microbial composition of enrichment cultures. (a) The relative abundance of microbial populations based on 16S rRNA gene-tag sequencing analysis before purification by adding antibiotics. (b) Growth curves of DL1YTT001 in anaerobic medium supplemented with lignin. (c) The relative abundance of microbial populations based on 16S rRNA gene-tag sequencing analysis after purification by adding antibiotics.

The strain DL1YTT001 was mesophilic with an optimum growth temperature of 20°C (Fig. S1). Hybridization Chain Reaction-Fluorescence In-Situ Hybridization (HCR-FISH), combined with scanning electron microscopy (SEM)/transmission electron microscopy (TEM) revealed that cells of this strain appeared as small cocci with a diameter of approximately 500 nm (Fig. S2). Cells occurred individually or in chain-like aggregates of several cells. Their size and morphology are similar to Bathyarchaeial cells previously observed from marine sediments by Catalyzed Reporter Deposition-Fluorescent In-Situ Hybridization (CARD-FISH)^20^.

To further enrich the Bathyarchaeial strain DL1YTT001, the culture was supplemented with various antibiotics (ampicillin, vancomycin, kanamycin, and streptomycin, each at a 50 μg/ml) (Fig. S3). The combination of antibiotics efficiently inhibited Bacterial growth, but also mildly suppressed the growth of strain DL1YTT001. This led to a hypothesis that the growth of strain DL1YTT001 might require some unknown substrates produced by the growing Bacteria, perhaps certain Bacterial metabolites. Subsequently, the culture was transferred to fresh media supplemented with the above four antibiotics as well as an additive of Bacterial metabolites prepared from supernatant from the same culture (Fig. S3). After three transfers, strain DL1YTT001 reached 89% of all microbes in the culture (Fig. 1c and Table S1), with a small fraction of Bacteria, including *Alkalibacteraceae (Firmicutes*) 2.54%, *Syntrophotaleaceae* (*Desulfobacterota*) 1.84%, *Spirochaetaceae* (*Spirochaetota*) 1.68%, *etc*.

Here we propose the candidate name ‘Ca. Marisediminiarchaeum ligniniphilus’ strain DL1YTT001 for this Bathyarchaeial strain. Etymology: Marisediminiarchaeum, Ma.ri.se.di.mi.ni.chae’um. L. neut. n.mare, the sea; L. neut. n. sedimen-inis, sediment; N.L. neut. n. archaeum, ancient one, archaeon; N.L. neut. n. Marisediminiarchaeum, an archaeon from the marine sediment. Ligniniphilus, lig.ni.ni’phi.lus. N.L. neut. n. ligninum, lignin; N.L. adj. philus-a-um,friend, loving; from Gr. adj. philos-ê -on, loving; N.L. masc. adj. ligniniphilus, lignin-loving, isolated as a lignin degrader with lignin as a single organic carbon source.

### Central carbon metabolism

A high-quality metagenomic assembled genome (MAG) of Bathyarchaeial strain DL1YTT001 was assembled from the enrichment culture metagenome with 99.22% completeness and 2.8% contamination (Table S2). Phylogenetic inferences based on the 16S rRNA gene and 37 concatenated Archaeal conserved proteins placed DL1YTT001 in subgroup-8 from a previous taxonomy for the phylum *Bathyarchaeota*^21^ (Fig. S4), and in the order-7 from a newly proposed taxonomy for the class *Bathyarchaeia*^22, 23^ (Fig. 2). For central carbon metabolism, DL1YTT001 contains a complete glycolysis/gluconeogenesis pathway, an Archaeal Wood-Ljungdahl (WL) pathway, an acetogenesis pathway, and a partial TCA cycle pathway and pentose phosphate pathway (Fig. 3 and S5, Table S3), similar to descriptions of genomes of most Bathyarchaeial groups^21, 22, 27^.

**Fig. 2.**
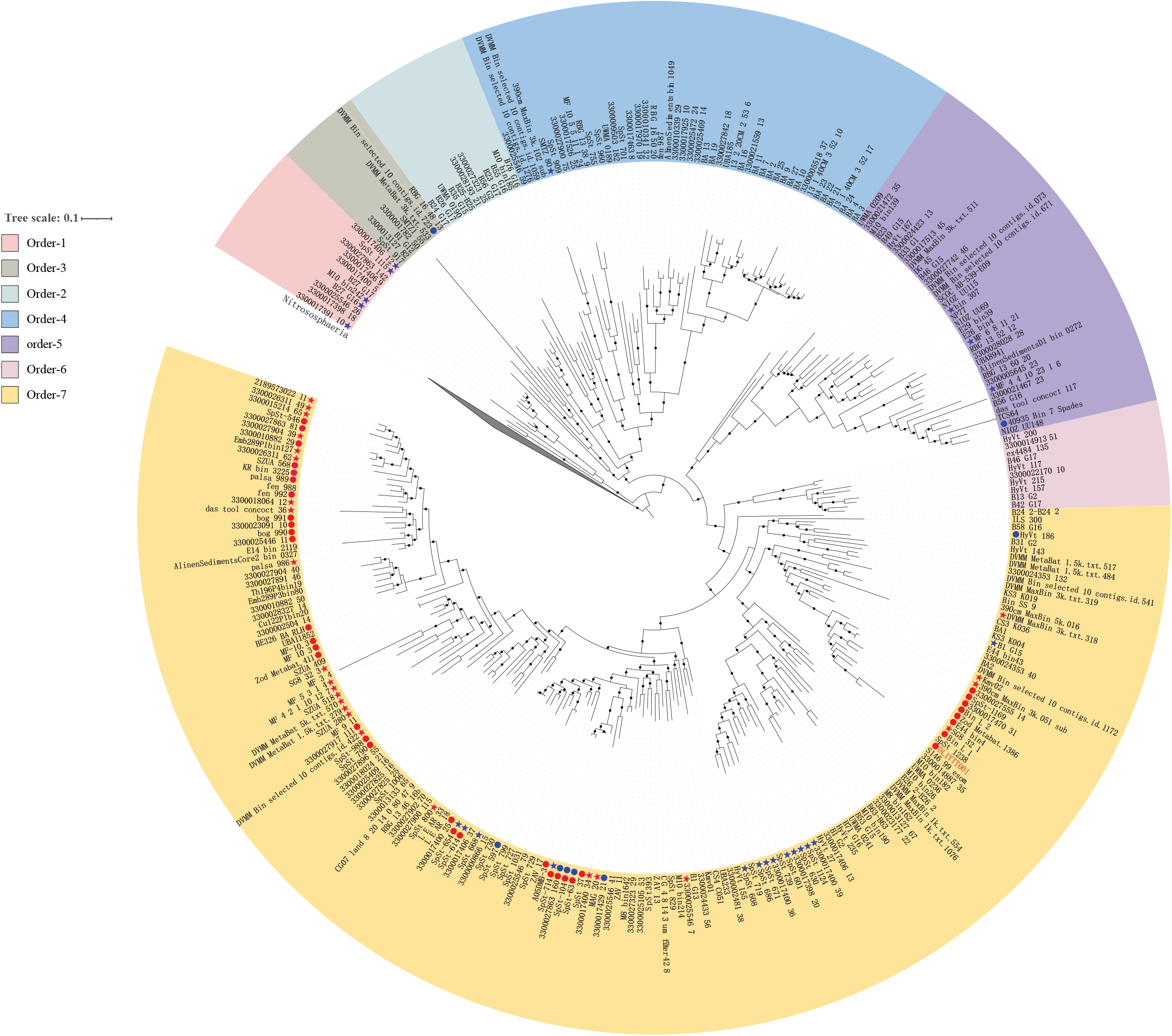
Maximum-likelihood phylogeny of *Bathyarchaeia* based on concatenated alignments constituting 37 concatenated ribosomal proteins. Bootstrap values were calculated from 1,000 iterations using fasttree. Bootstrap values of >70% are labeled with black dots. MAGs containing *Bathyarchaeia* specific MtgB genes and gene clusters are highlighted with star- and dot-shaped red labels, respectively. MAGs containing Bacterial-type MT1 genes and gene clusters are highlighted with star- and dot-shaped blue labels, respectively.

**Fig. 3.**
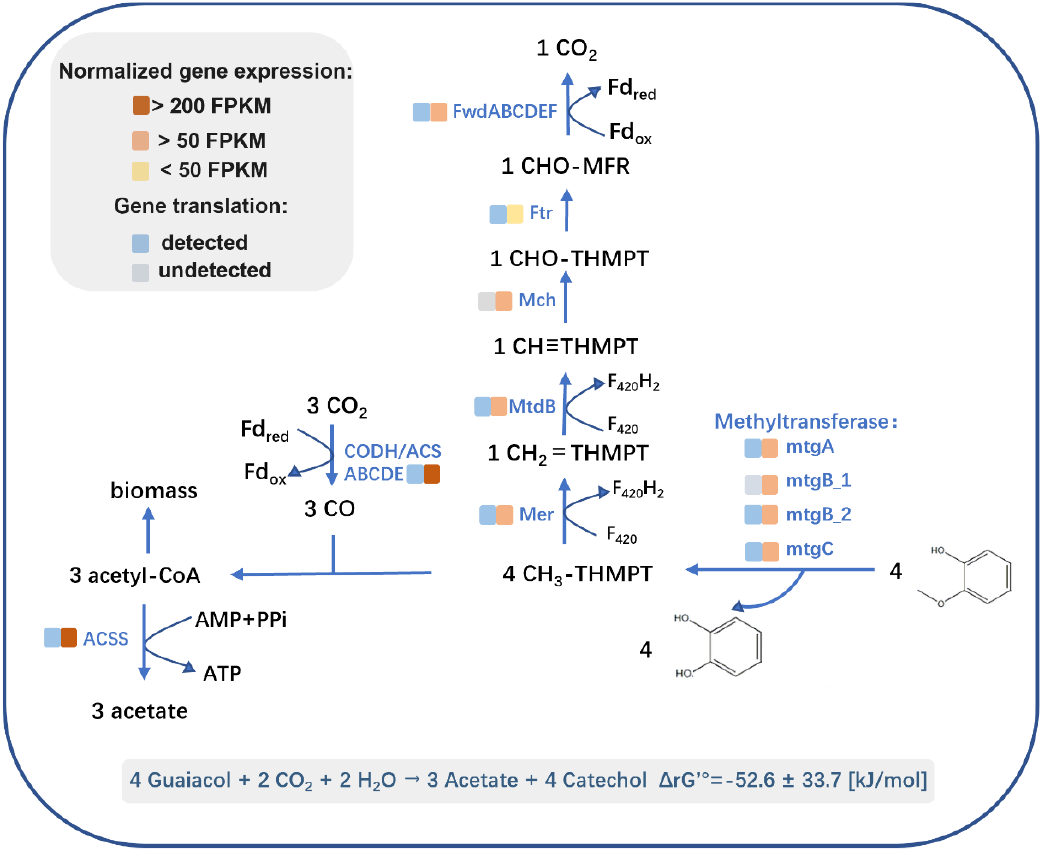
An overview of the central metabolic pathway of DL1YTT001. The process of O-demethylation of guaiacol is accompanied by the fixation of carbon dioxide and the production of acetate. Yellow rectangles of varied shades exhibit different gene transcription levels, and blue rectangles of different shades show whether the relevant proteins were detected in the proteomics. Below the pathway is the Gibbs free energy (ΔG) estimated for the overall equation.

Gene transcription and translation patterns of strain DL1YTT001 were also analyzed. In the central carbon metabolism, acetogenesis through the WL pathway, considered an essential metabolic process for *Bathyarchaeia*^18,26^, showed high transcription in the transcriptome, and most of the associated proteins were detected in the proteome (Fig. 3 and S5, Table S3). Acetyl-CoA synthetase (Acss) and monoxide dehydrogenase/acetyl-CoA synthase complex (Codh/Acs), key enzymes of acetogenesis and the WL pathway, respectively, showed high transcript levels with FPKM (fragments per kilobase of transcript per million mapped reads) values > 200. Within the acetogenesis pathway, only one enzyme N5,N10-methenyltetrahydromethanopterin cyclohydrolase (Mch) was absent in the proteome, likely due to the low biomass of strain DL1YTT001 that was inadequate to produce a detectable level of this enzyme.

Considering that strain DL1YTT001 was grown in a medium in which lignin was the sole organic carbon source, we searched for genes related to lignin degradation in its MAG. Still, no known genes coding for lignin polymer depolymerization enzymes were found. We attempted to verify that the mode of lignin metabolism of strain DL1YTT001 is similar to that described for acetogenic Bacteria with the capability of O-demethylation from lignin-derived methoxylated aromatic compounds (ArOCH_3_) instead of polymer depolymerization^26^. Known Bacterial-type methyltransferase systems previously identified both in Archaea and Bacteria ^16, 17^were searched but were also absent in the DL1YTT001 MAG. We speculated that this strain might contain a novel transmethylation system different from those known Bacterial-type systems. To verify that O-demethylation occurs during the incubation of DL1YTT001 and check whether there is a novel O-demethylation pathway, the metabolome targeting low molecular weight aromatic compounds was analyzed along with transcriptomic and proteomic data.

### O-demethylation from lignin-derived ArOCH_3_ occurring during incubation

The metabolome targeting low molecular weight aromatic compounds on day 0 and day 30 of the incubation was analyzed by GC-MS (Table 1). On day 0, a variety of ArOCH_3_ as lignin-derived monomers was detected with varied concentrations, including guaiacol (0.8-1.0 μg/ml), vanillin (2.5-7.3 μg/ml), acetovanillone (6.2-7.4 μg/ml), vanillic Acid (11.2-12.9 μg/ml), homovanillic acid (2.7-3.0 μg/ml) and syringic acid (1.4-1.5 μg/ml). These monomers all decreased dramatically after 30 days of incubation, with a substantial accumulation of their demethylated products, including catechol and protocatechoic acid. Thus, O-demethylation from these ArOCH_3_ occurred during the incubation of strain DL1YTT001.

**Table 1.** GC-MS analysis of low molecular weight aromatic compounds in the DL1YTT001 culture sampled at 0 day and 30 days of the incubation. All assays were performed in duplicate. Media without inoculation of the DL1YTT001 culture were used as negative controls.

| Retention<br>(min) | Compound (µg/ml) | DL1YTT001-1 |  | DL1YTT001-2 |  | medium-1 |  | mediun-2 |  |
| --- | --- | --- | --- | --- | --- | --- | --- | --- | --- |
|  |  | 0 day | 30 days | 0 day | 30 days | 0 day | 30 days | 0 day | 30 days |
| 11.6 | Guaiacol | 1.0 | 0.0 | 0.8 | 0.0 | 0.9 | 0.8 | 1.0 | 1.1 |
| 18.29 | Vanillin | 6.6 | 0.2 | 2.5 | 0.1 | 5.1 | 4.7 | 7.3 | 6.4 |
| 19.92 | Acetovanillone | 6.9 | 0.0 | 6.2 | 0.0 | 7.0 | 6.9 | 7.4 | 7.0 |
| 22.61 | Vanillic Acid | 12.6 | 0.0 | 11.2 | 0.0 | 12.1 | 12.8 | 12.9 | 12.8 |
| 22.68 | Homovanillic Acid | 2.9 | 1.7 | 2.7 | 0.2 | 2.9 | 3.0 | 3.0 | 3.0 |
| 24.82 | Syringic acid | 1.4 | 1.4 | 1.4 | 0.0 | 1.5 | 1.5 | 1.4 | 1.5 |
| 13.62 | Catechol | 0.0 | 6.1 | 0.0 | 7.1 | 0.0 | 0.0 | 0.0 | 0.0 |
| 23.52 | Protocatechoic acid | 0.0 | 5.0 | 0.0 | 0.59 | 0.0 | 0.0 | 0.0 | 0.0 |

### The novel O-demethylation pathway involves methyl transfer from ArOCH_3_ to tetrahydromethanopterin (H_4_MPT)

Although there is no known Bacterial-type methyltransferase gene cluster in the MAG of DL1YTT001, according to the compositional gene order of these gene clusters and comprehensive analysis of genomic, transcriptomic, and proteomic data, we found a novel gene cluster from the DL1YTT001 MAG that may carry out the O-demethylation/methyltransferase process on ArOCH_3_. This putative methyltransferase complex includes two copies of methyltransferase 1 (MT1) (MtgB_1 and MtgB_2), one copy of methyltransferase 2 (MT2) (MtgA), and one copy of corrinoid protein (CP, MtgC) (Fig. 4a). The genes within the DL1YTT001 methyltransferase complex were all transcribed at high levels as indicated by FPKM expression values, and proteins of MtgA, MtgB_2, and MtgC were also detected in the proteome (Fig. 3 and S5, Table S3). Within these complexes, MtgB_1 and MtgB_2 are proposed to catalyze the demethylation of ArOCH_3_ and transfer the resulting methyl group to MtgC, and MtgA is proposed to catalyze a subsequent transfer of the methyl group from MtgC to H_4_MPT (Fig. 4b). The gene order of the methyltransferase complex in the MAG of strain DL1YTT001 is also similar to the methyltransferase complexes of methanol, methylamine (monomethylamine, dimethylamine, and trimethylamine), glycine, betaine, etc (Fig. 4c).

**Fig. 4.**
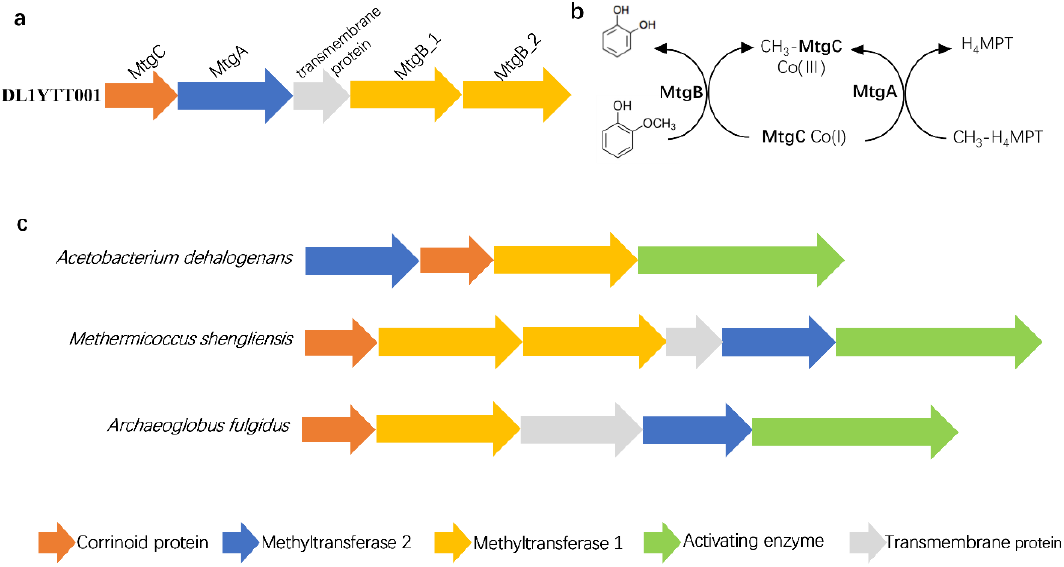
The operon encoding the gene cluster of O-demethylase/methyltransferase complexes. The O-demethylase/methyltransferase gene cluster (a) and the methyl-transfer pathway (b) of DL1YTT001 with guaiacol as the substrate; (c) shows the gene clusters encoding *Acetobacterium dehalogenans* vanillate O-demethylase system^14^, *Methermicoccus shengliensis* methoxybenzoate O-demethylase system^16^, and *Archaeoglobus fulgidus* 2-methoxyphenol O-demethylase system^17^. Co(I) and Co(III), corrinoid protein with cobalt in their respective valence states; H_4_MPT, tetrahydromethanopterin.

In the methyltransferase complex, MT1 is the key enzyme responsible for substrate selectivity and determines the type of methyl substrate. Neither of the two putative MT1s (MtgB_1 and MtgB_2) of strain DL1YTT001 have homology with known Bacterial or Archaeal sequences. Additionally, both of the protein sequences had relatively short lengths of only ~300 amino acids (~32kDa), compared with known MT1 lineages whose lengths were reported to be ~400 amino acids (~48kDa) (Fig. 4). All evidence indicates that these two MT1s (MtgB_1 and MtgB_2) are novel.

MT2 and CP, which are not involved in methyl substrate specificity, show homology with various other methyl substrate methyltransferase systems, unlike MT1. In the DL1YTT001 MAG, the genes of CP (MtgC) and MT2 (MtgA) also show homology with the known CP and MT2 lineages from various methyl substrate methyltransferase systems such as *Methermicoccus shengliensis* methoxybenzoate ArOCH_3_ O-demethylase system^16^, *Acetobacterium dehalogenans* vanillate ArOCH_3_ O-demethylase system^14^, and *Desufitobacterium hafniense* glycine betaine demethylase system ^28^(Fig. S6). DL1YTT001 likely acquired the CP (MtgC) and MT2 (MtgA) for ArOCH_3_ metabolism from other Bacteria or Archaea through horizontal gene transfer.

In addition to MT1, MT2, and CP, some reported methyltransferase systems also include an activating enzyme (AE) for the reductive activation of corrinoid, which was apparently absent or unidentified in the DL1YTT001 genome (Fig. 4a). As is the case with MtgB_1 and MtgB_2, the unidentified AE in DL1YTT001 is likely also novel and would require further identification. In addition, because these methyltransferase systems function in the cytoplasm, their cells require transporters for the uptake of ArOCH_3_. A membrane protein adjacent to the DL1YTT001 methyltransferase complex was identified in the DL1YTT001 genome and considered a candidate for the transport protein for aromatic compounds (Fig. 4a).

### Enzyme activity of novel methyltransferase 1 (MtgB_2)

To verify the function of the novel MT1s of DL1YTT001 in O-demethylation and methyl transfer, the MtgB_2 (which was detected in the proteome) and MtgC of DL1YTT001 were expressed and purified in *Escherichia coli* and subjected to an enzyme activity assay (Fig. S7). In addition, for the synthesis of the Co(I) state of CP, which is the active substrate of MT1 (MtgB_2), an AE gene from the acetogenic Bacteria *Acetobacterium dehalogenans* was also expressed and purified in *E. coli* and included in the enzymic reaction. Firstly, AE reactivated the Co(II) state of MtgC by reducing the cobalamin to the active Co(I) state with the use of ATP and titanium(III) citrate, causing a blue shift in the absorption peak from 480 nm to 390 nm, as shown in the UV-vis spectrum (Fig. 5). Subsequently, the addition of methoxylated substrate (guaiacol) and the MtgB_2 resulted in a red shift in the absorption peak from 390 nm to 520 nm, suggesting the formation of methyl-Co-(III), i.e., the occurrence of methyl transfer. The HPLC chromatogram results show a decrease in the guaiacol peak along with an increase in the catechol peak, also suggesting the occurrence of O-demethylation (Fig. S8). All these results confirmed the *in vitro* activity of the novel MtgB_2, which was expressed during the growth of DL1YTT001 cells and played a part in transferring methyl groups of ArOCH_3_.

**Fig. 5.**
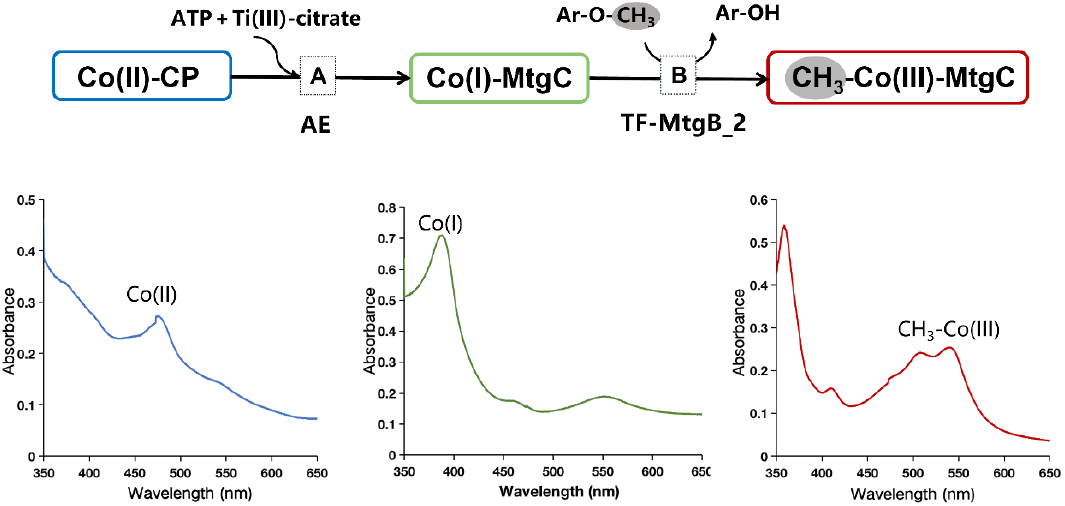
The DL1YTT001 O-demethylation/methyltransferase activities with guaiacol as the substrate. Reaction A: Activating enzyme (AE), ATP, and titanium (III) citrate are required for the activation of corrinoid protein (MtgC) from the Co(II) state (blue) to the active Co(I) state (dark green); AE was from *Acetobacterium dehalogenans* DSM 11527 (GenBank accession no. ACJ01666.1). Reaction B: MtgB transfers the methyl group from guaiacol to Co(I)-MtgC, resulting in methylated Co(III)-MtgC (red). Conversion of guaiacol to catechol was confirmed by HPLC (Fig. S6). All bottom panels correspond to UV/visible spectra measured after each reaction reflecting the different states of the cobalamin carried by MtgC.

## Discussion

### Growth of DL1YTT001 depends on as yet unidentified metabolites produced by co-cultured Bacteria

A substantial fraction of organic carbon is buried in nearshore sediments where the microbial transformation of organic carbon occurs as a key process influencing carbon flow and ultimately atmospheric oxygen and carbon dioxide concentrations^29^. Nevertheless, many questions about how microorganisms regulate organic carbon cycling remain unanswered, in particular the biogeochemical role of Archaea, which are often abundantly distributed in nearshore sediments^30–32^. *Bathyarchaeia* are one of the predominant Archaeal groups in global nearshore sediments but to date are poorly characterized due to challenges with isolation. Members of *Bathyarchaeia* have been proposed to have significant ecological impacts on global carbon cycling based on genomic analyses^18, 19, 21, 22, 24^, stressing the need for increased cultivation efforts for this Archaeal group. From a yearlong lignin enrichment of intertidal sediments in 2018^26^ and five years of continuous transfers, we obtained a 60% enrichment culture of Bathyarchaeial strain DL1YTT001 (Fig. 1). The growth of strain DL1YTT001 was inhibited by Bacteria-targeted antibiotics. However, following the addition of a metabolite-rich culture supernatant, DL1YTT001 resumed growth (to 89% relative abundance) (Fig 1 and Fig S3), indicating that growth of DL1YTT001 depends on as yet unidentified metabolites produced by co-cultured bacteria in the enrichment. This dependence is shown by the inability of DL1YTT001 to grow in the absence of some essential cofactors and the genome’s lack of genes for their synthesis. Some genes are missing in certain biosynthetic pathways, such as most of the genes for coenzyme F_420_ biosynthesis (Fig S5). Because individual species often lack the machinery to produce all essential factors necessary for growth and survival, interspecies exchange of secondary metabolites, growth factors, signaling molecules, and even electrons is crucial within microbial communities^33^. This is often the case with Archaea that require a syntrophic partner, such as the recently cultured Lokiarchaeia archaeon MK-D1 that requires interspecies transfer of amino acids and vitamins from a methanogen partner and anaerobic methane-oxidizing Archaea (ANME) that transfers electrons to a sulfate-reducing partner to facilitate cooperative growth^34, 35^. The mechanisms of interaction between strain DL1YTT001 and potential syntrophic partners will be explored in further physiological experiments. This includes the determination of potential interspecies factors, which may help design an ideal culture medium supplemented with these factors to facilitate the isolation of a pure culture of DL1YTT001 without any co-cultured partners.

### *Bathyarchaeia*-specific methyltransferase systems likely constitutes a vital step in the anaerobic degradation of lignin

Anaerobic biodegradation of lignin in marine sediments, particularly in nearshore sediments, is likely a globally important microbial process and significant for the cycling of organic carbon. O-demethylation from lignin-derived methoxylated aromatic compounds (ArOCH_3_) is an important step of lignin biodegradation likely performed by acetogenic Bacteria and several lineages of Archaea^17^. These microorganisms acquired and shared Bacterial-type methyltransferase systems by horizontal gene transfer^17^. Recently, one MAG of *Bathyarchaeia* has also been reported to carry a Bacterial-type methyltransferase system^17^. In this study, the gene distribution of Bacterial-type methyltransferase in 298 MAGs of *Bathyarchaeia* were further analyzed, and we found that 31 MAGs contain a Bacterial-type methyltransferase gene cluster or the gene of Bacterial-type methyltransferase 1 (MT1); these MAGs are distributed in five of the seven orders of *Bathyarchaeia* (Fig. 2 and Table S4). Interestingly, 80% of these 31 MAGs containing Bacterial-type MT1 were assembled from high-temperature environments such as hydrothermal or hot spring sediments. This suggests that horizontal transfer of Bacterial-type methyltransferase systems may confer an adaptive advantage for some members of *Bathyarchaeia* in high-temperature environments.

A process of O-demethylation from lignin-derived ArOCH_3_ occurred in the culture system of Bathyarchaeial strain DL1YTT001 (Table 1). However, a Bacterial-type methyltransferase system is absent in the DL1YTT001 genome, so there likely exists a novel methyltransferase system. And indeed, within the genome, we found two MT1 genes (MtgB_1 and MtgB_2) that are used for substrate selection and methyl transfer from ArOCH_3_ to cobalamin, both of which are distinct from Bacterial-type MT1 in sequence identity and length. The two copies of MT1 (MtgB_1 and MtgB_2) share a sequence similarity of 44.85%. This dissimilarity suggests different substrate affinities for the two proteins, which may explain why the DL1YTT001 cultures could demethylate various types of ArOCH_3_ (Table 1). Public genomic data was searched extensively for genes coding for homologous enzymes of MtgB. We found that unlike the genes of Bacterial-type MT1, which are widely detected in both Bacterial and Archaeal lineages, the genes encoding MtgB were only found in genomes of *Bathyarchaeia*. In 298 Bathyarchaeial MAGs, 60 MAGs were found to contain the *Bathyarchaeia*-specific MtgB or its gene cluster, and all 60 MAGs clustered in order-7 (Fig. 2 and Table S4). Thus, the evolution model of *Bathyarchaeia-specific* MtgB gene may be vertical inheritance, and the original host of this gene remains to be discovered. These homologous enzymes of the MtgB clustered into three groups based on phylogenetic analysis (Fig. S9). The MtgB_1 and MtgB_2 of DL1YTT001 clustered within group-1 and group-2, respectively. Since group-3 showed a long evolutionary distance from the former two groups, it is unclear whether group 3 would retain the same function. *Bathyarchaeia*-specific MtgB may be more efficient compared to Bacterial-type MT1, resulting in the Bathyarchaeial strain DL1YTT001 outcompeting Bacteria and becoming the most abundant microorganism in the lignin culture system (Fig. 1a).

DL1YTT001 belongs to subgroup Bathy-8 of order-7 *Bathyarchaeia* (Fig. 2 and S4)^22^. Bathy-8 is frequently detected in high abundances in ocean-margin sediments, especially in deeper sediment layers with low redox potential and lacking electron acceptors^20, 25, 36–38^. Thus, O-demethylation from ArOCH_3_ catalyzed by these *Bathyarchaeia*-specific methyltransferase systems likely constitutes a vital step in the anaerobic degradation of lignin in ocean margin sediments. Order-7 is the biggest order of *Bathyarchaeia* and is phylogenetically diverse (Fig. 2)^22^. More than a third of MAGs in order-7 contained MtgB or its gene cluster; these MAGs were assembled from diverse anaerobic habitats, including termite guts, mangrove sediments, subsurface fracture fluids, permafrost active layer soils, marine sediments, peatlands, peat soils, estuary sediments, hot spring sediments, and mud volcanos (Table S4). Order-7 may play an important role in O-demethylation from ArOCH_3_ in these anaerobic habitats, and MtgB may help order-7 adapt to diverse anaerobic environments, allowing this lineage to become ubiquitous. Furthermore, order-7 is not a deep-branching clade in the phylogenetic tree of class *Bathyarchaeia* (Fig. 2), implying a late origin of these *Bathyarchaeia-specific* methyltransferase genes. Considering the potential role of these enzymes in lignin degradation, they may have evolved after the emergence of vascular plants in the mid-Cambrian to early Ordovician^39, 40^. This hypothesis needs to be confirmed by further evolutionary analyses.

### Central carbon metabolism involves the oxidation of methyl groups and reduction of CO_2_

In addition, for the central carbon metabolism of *Bathyarchaeia*, we reported earlier that Bathy-8 (a predecessor culture of DL1YTT001) incorporated ^13^C labeled inorganic carbon (IC) for biomass synthesis and acetate production with lignin as an energy source^26^. In this study, during the growth of strain DL1YTT001, all the genes in the pathway of acetogenesis (acetate production involves a CO_2_ reduction step via the WL pathway) were highly transcribed, and most of their proteins were present in the proteome (Fig. 3 and S5, Table S3). Meanwhile, the *Bathyarchaeia*-specific methyltransferase system was also transcribed and translated. Thus, the ArOCH_3_-derived methyl carried by H_4_MPT would be oxidized to CO_2_; this process would result in the accumulation of reducing equivalents distributed among multiple electron carriers (e.g., in the forms of reduced ferredoxin and F_420_H_2_). Strain DL1YTT001 was also capable of converting the methyl group carried by H4MPT to acetyl-coenzyme A (CoA) by addition of CO_2_ which would consume these reducing equivalents. DL1YTT001 may be able to alternate between oxidative and reductive metabolism through accumulation and consumption of cellular reducing equivalents. Furthermore, the produced acetyl-CoA can be further used for biosynthesis and acetate production, where ATP can be directly generated from acetyl-CoA conversion to acetate. Such recycling of reducing equivalents has also been observed in acetogenic Bacteria and methanogenic Archaea that conduct CO_2_-reducing acetogenesis/methanogenesis which allows cells to re-oxidize excess reducing power produced from oxidation of ArOCH_3_-derived methyl groups^16, 41, 42^ In the case of DL1YTT001, after transfer from ArOCH_3_ to H4MPT via the methyltransferase system, one-quarter of the methyl groups were oxidized to CO_2_, while the rest was converted to acetyl-CoA for acetate production (Fig. 3). With guaiacol as the substrate, the overall reaction yields a Gibbs free energy of −52.6±33.7 kJ/mol (Fig. 3). This low energy yield may be the reason for its slow growth (Fig 1b).

## Conclusion

In conclusion, we have achieved a continuous laboratory culture of the ubiquitous *Bathyarchaeia* from nearshore sediments. This Bathyarchaeial archaeon employs a methyltransferase system for transferring methyl from lignin-derived ArOCH_3_. The system is specific to *Bathyarchaeia* and particularly prevails in order-7. Given the high abundance of *Bathyarchaeia* and lignin-derived organic carbon in nearshore sediments, demethylation catalyzed by this *Bathyarchaeia*-specific methyltransferase system is likely a critical step within the anaerobic lignin degradation pathway in these environments. Furthermore, we found that the carbon central metabolism of this Bathyarchaeial archaeon involves the oxidation of methyl groups, reduction of CO_2_, and production of acetate. There are still many questions that need to be answered in future work, e.g., whether the members of *Bathyarchaeia* can further crack the hydroxylated derivatives (ArOH) after demethylation of ArOCH_3_, and whether they participate in the depolymerization of lignin polymers. The Bathyarchaeial enrichments we have obtained will facilitate in-depth investigation into these questions, enabling us to better understand the ecological significance of the *Bathyarchaeia*.

## Materials and Methods

### Sample Collection

Sediment samples were collected from an intertidal zone (30.592817 N, 122.083493 E) on Dayangshan island in Hangzhou Bay of the East China Sea^26^. Samples were kept in oxygen-free gas-tight bags on ice, transported to the laboratory within 3 hours, and then stored at 4°C until further culture experiments.

### Cultivation Conditions

Initial cultivation of the intertidal sediments has been described previously^26^. Routine transfers were performed at 2-3 months intervals using a sulfate-free artificial seawater medium supplemented with 50 mM NaHCO3 and 5 g/L alkali lignin (Sigma, CAS:8068-05-1). For nucleic acid and protein extraction and chemical parameter measurements, 2 mL/10 mL of each culture was collected and centrifuged at 13,800 ×g for 10 min to separate supernatants from pellets then stored at −80°C until analysis.

### DNA and RNA extraction, metagenome and metatranscriptome sequencing

DNA extraction for 16S rRNA gene amplification was performed using the PowerSoil DNA Isolation Kit (Qiagen). Total DNA for metagenome sequencing was obtained using an Advanced Soil DNA Kit (mCHIP). Total RNA was recovered from one-month-old of the DL1YTT001 culture by using a RNeasy^®^ PowerSoil^®^ kit (Qiagen). The kits mentioned above were used according to manufacturer protocols.

Libraries for metagenome sequencing were generated using NEB Next^®^ Ultra™ DNA Library Prep Kit for Illumina^®^ (New England Biolabs), following the manufacturer’s recommendations with the addition of index codes. For metatranscriptome sequencing, the detailed methods involving the removal of ribosomal RNA (rRNA) and libraries construction are described in the Supplementary Methods. The metagenomic and metatranscriptomic libraries were sequenced on an Illumina Novaseq6000 platform (Illumina, USA), generating 150 bp paired-end reads.

### Q-PCR

Bacterial and bathyarchaeotal 16S rRNA genes were quantified by qPCR and were amplified using the primers Bac341F/prokaryotic519R^43^ and Bathy-442F/Bathy-644R^38^, respectively. The reaction mixture included 10 μL of SYBR Premix Ex Taq (TaKaRa), 0.4 μL of ROX reference dye (50×; TaKaRa), 0.8 μM of each primer, and 1 μL of template DNA. Clones with Bacterial and bathyarchaeotal 16S rRNA genes^38^ were used for standard curve construction. The abundance of each targeted gene in the DNA assemblage was determined in triplicate analyses.

### 16S rRNA gene amplification and analysis

The hypervariable V4 region of the prokaryotic 16S rRNA genes was amplified using primer sets 515F-806R^44^. The thermal cycling program was as follows: initial denaturation at 95°C for 4 minutes, 30 cycles at 95°C for 30 seconds, 50°C for 60 seconds, 72°C for 60 seconds, and a final extension at 72°C for 7 minutes. Each reaction (50 μL) contained 10×PCR buffer, dNTPs (100 μM each), 0.25 μM of each primer, 2.5 U of DNA polymerase (Ex-Taq; TaKaRa, Dalian, China), and approximately 10 ng of total DNA. The PCR products were purified using an E.Z.N.A. Gel Extraction Kit (Omega Bio-Tek, Norcross, GA, USA) according to the manufacturer’s instructions. Sequence reads were obtained from the Illumina NovaSeq platform based on 2 × 250 bp cycles and the NovaSeq 6000 SP Reagent Kit (500 cycles, Illumina, San Diego CA, USA). Further analysis was performed using the QIIME 2 standard pipeline (Version 2020.11)^45^.

### Metagenomic assembly, binning, and annotation

Trimmomatic was used to remove possible adaptors and low-quality bases for each read^46^. Trimmed and paired-end DNA reads were assembled using SPAdes De Novo Assembler^47^. For assembled contigs longer than 1 Kb, open reading frames (ORFs) were predicted and translated using prodigal (v2.6.3) with -p meta parameters^48^. Assembled contigs (longer than 1 kb) were binned into putative taxonomic groups based on abundance information using MaxBin version 2.2.4 with the run MaxBin.pl script^49^. Detailed estimates of genome contamination and completeness were assessed based on lineage-specific marker sets with CheckM^50^. The retrieved MAGs were annotated using eggnog-mapper-1.03 in the EggNOG database with an e-value of 10_-_^10^.The completeness of specific pathways and functions within the MAGs was assessed based on the canonical pathways available in KEGG Pathway Database (www.kegg.jp).

### Metatranscriptomic analysis

For metatranscriptomic analysis, sequence reads were filtered to remove low-quality and rRNA reads, then mapped to metagenome assembled contigs with TopHat v2.1.1^51^. FPKM (expected fragments per kilobase of transcript per million fragments mapped) values were used to estimate the expression level of each gene using Cufflinks 2.2.1 scripts (http://cole-trapnell-lab.github.io/cufflinks/).

### HCR-FISH

HCR-FISH was carried out as previously described^52^. A universal Bacteria-specific probe EUB338^53^ and Bathyarchaeia-specific probe Bathy-442^38^ were used in this study.

### Proteomic Analysis

After seven days of cultivation, cell pellets were harvested under anaerobic conditions from 10 ml of DL1YTT001 culture by centrifugation at 13,000 × g for 25 min, frozen in liquid nitrogen, and stored at −80 °C. The protein of cell pellets was extracted for LC-MS/MS analysis using an Easy nLC1200/Q Exactive plus mass spectrometer, the detailed sample preparation and analysis method using LC-MS/MS is described in the Supplementary Methods.

### GC-MS analysis of low molecular weight aromatic compounds

GC-MS was used to monitor low molecular weight aromatic compounds across the incubation process. Basic experimental procedures were carried out following the methodology of Raj et al.^54^, with some modifications. Specifically, aliquots (2 mL) of enrichment cultures or controls were sampled at 0 d and 30 d, low-molecular-weight (LMW) aromatic compounds were extracted, then identified and quantified on a Trace 1310 gas chromatograph coupled to a TSQ8000 mass spectrometer (Thermo Fisher Scientific, USA) using a HP-5MS capillary column (30 m × 0.25 mm i.d., 0.25 μm film thickness). The detailed sample preparation and analysis method using GC-MS is described in the Supplementary Methods.

### The heterologous protein production of MtgC and MtgB

The genes encoding the methyltransferase I (MtgB_2) and corrinoid protein (MtgC) were amplified from the DL1YTT001 culture for cloning in expression vector pCold-TF and pET-28a, respectively. For Co(I) production, the activating enzyme (AE) gene of Acetobacterium dehalogenans DSM 11527 (GenBank accession no. ACJ01666.1) was synthesized, then amplified for cloning in expression vector pET-28a. For the production of MtgB_2, MtgC, and AE, the plasmids were used for transformation into *E. coli* BL21 (DE3). The detailed method of heterologous protein production is described in the Supplementary Methods.

### Enzyme activity assays

The protocol for enzyme activity assays were adapted from Schilhabel et al.^14^ and Kurth et al.^16^, which performed in 350 μl anaerobic Quartz cuvettes (Purshee Co., Ltd, Yixing, China), which were sealed with rubber stoppers and gassed with N_2_. MtgB protein activity was determined in a total 300 μl buffer containing 35 mM Tris HCl (pH 7.5) and 70 mM KCl. Firstly, reconstituted Co(II)-MtgC at 1.2 mg/ml (~79 μM) final concentration was activated by adding 12 mM MgCl_2_, 0.5 mM Ti(III) citrate (freshly prepared), and 2.3 mM ATP. The conversion to Co(I)-MtgC was followed by a change in absorbance at 387 nm on a HACH device (spectrophotometer DR/5000 Company, USA). The enzymatic reaction was started by the addition of 2.3 mM guaiacol and MtgB at a final concentration of 0.015 mg/ml. Fifty microliters of the sample were collected before the addition of MtgB protein and after the activity assay for analysis of methoxy aromatic compounds and products by HPLC (see below). Meanwhile, the formation of CH3-Co(III)-MtgC from Co(I)-MtgC was expected to result in a decrease in absorption at 387 nm and an increase in absorption at 530 nm on a spectrophotometer. As a negative control, 2.3 mM methanol was used.

For determination of the occurrence of bioconversion catalyzed by the MtgB protein, substrates and products of the biochemical reaction were directly analyzed on an Agilent 1260 Infinity HPLC system equipped with UV detector and ZORBAX Eclipse Plus C18 column (100×4.6mm, particle size 3.5 mm)^14^. H_2_O/MeCN (containing 0.1% TFA) was used as the eluent at a flow rate of 1 mL/min. A linear gradient was applied: 100% H_2_O (hold 2 min), gradient from 0-40% MeCN (13min), gradient to 100% MeCN (5 min, hold 2 min), 100% H_2_O (hold 3min). Compounds were detected by UV-absorption, and peaks were identified by co-injection of commercially bought reference material.

## Acknowledgements

We thank Jicheng Yao (Hohai University), Jialin Hou (Shanghai Jiao Tong University) and Ruize Xie (Shanghai Jiao Tong University) for help with the metagenome analysis, Xipeng Liu (Shanghai Jiao Tong University) and Qinying Huang (Shanghai Jiao Tong University) for their kind suggestions for the enzyme activity test. This work was supported financially by the Natural Science Foundation of China (Grants 42276139, 41867057, 42141003, 41921006, 92051116, 91951209) and National Postdoctoral Program for Innovative Talents (Grant No. BX20190204).

## Authors’contributions

TY and FW designed research; TY, HH and XZ performed research; TY analyzed data; and TY, HH, YW, DP and FW wrote the paper.

