## supplementary for "Cultivation of widespread *Bathyarchaeia* reveals a novel methyltransferase system utilizing lignin-derived aromatics"

### Supplementary Methods

#### Removal of ribosomal RNA (rRNA) and library construction for metatranscriptome sequencing

For metatranscriptome sequencing, Bacterial and Archaeal 16S and 23S ribosomal RNA (rRNA) transcripts in total RNA samples were reduced using the Ribo-off rRNA Depletion Kit (Bacteria) (Vazyme). Whole mRNAseq libraries were generated using NEB Next® Ultra™ Nondirectional RNA Library Prep Kit for Illumina® (New England Biolabs). Fragmentation was carried out using NEB Next First Strand Synthesis Reaction Buffer. First strand cDNA was synthesized using random hexamer primers and M-MuLV Reverse Transcriptase (RNase H), and second strand cDNA synthesis was performed using DNA Polymerase I and RNase H. After adenylation of 3’ ends of the DNA fragments, NEB Next Adaptor with hairpin loop structure was ligated to prepare for hybridization. cDNA fragments of 150~200 bp in length were selected with SpeedBead Magnetic Carboxylate Modified Particles (Global Life Sciences Solutions Operations,Buckinghamshire). PCR was performed with Phusion High-Fidelity DNA polymerase, Universal PCR primers and Index (X) Primer. At last, the PCR products were purified with AMPure XP beads, and the library insert size was assessed on the Qsep400 High-Throughput Nucleic Acid Protein Analysis system (Houze Biological Technology Co).

#### Sample preparation and analysis of proteome

Cell pellets were resuspended in 600 µL Lysis Buffer (8 M urea, 1% SDS, 100 mM $NH_4HCO_3$) in a 2 mL shock resistant tube with addition of 200 mg glass beads, followed by bead beating using a tissue lyser (Tissuelyser-48, Shanghai Jingxin, China) (two cycles at 30 Hz for 30 s with a 120 s interval at 4 °C). After centrifugation of the lysate at 14,000 × g for 30 min at 4 °C, the supernatants were transferred to a 1.5 mL Eppendorf tube. 6 µL DTT (1 M) was added to the tube, followed by incubation at 37 °C for 30 min. Proteins were then alkylated with 30 µL iodoacetamide (1M) and incubated at 37 °C for 20 min in the dark. Subsequently, proteins were precipitated with five volumes of cold acetone at -20°C overnight, harvested by centrifugation at 14,000

× g for 30 min at 4 °C, and air-dried at room temperature. Digestion was performed using 3 μg trypsin, followed by incubation at 56 °C for 2 h. The supernatants were collected by centrifugation at 14,000 × g for 30 min at 4 °C, desalted using Monospin C18 care (GL Sciences) and Pierce C18 spin tips (Thermo Scientific), and subsequently lyophilized in a SpeedVac centrifuge. The lyophilizate was reconstituted in 1% (v/v) formic acid prior to LC-MS/MS analysis. LC-MS/MS analysis of the samples was performed using an Easy nLC1200/Q Exactive Plus mass spectrometer. Peptides were loaded onto an Acclaim PepMap 100 (100 μm × 2 cm, NanoViper, C18, 5 μm, 100 A) (Thermo Scientific) and separated on an analytical column at a flow rate of 300 nL/min using a 120 min linear gradient, ranging from 0 to 80% of acetonitrile in a mobile phase.

The mass spectrometer was operated in positive mode only using fragmenting precursors with an assigned charge of ≥2. An isolation window of 1.2 m/z was used, and survey scans were acquired at 200-2000 m/z with resolution set to 70,000 at m/z 200. Fragmentation spectra were captured at 17,500 at m/z 200. Maximum ion injection time was set to 150 ms and 100 ms for MS and MS/MS scans, respectively, and dynamic exclusion of 30 s was applied.

The resulting MS/MS raw data was processed by proteome discoverer 2.4 (Thermo Scientific) with carbamidomethylation as a fixed modification and methionine oxidation as a variable modification. Data was filtered using a cutoff of 1% for peptide- and protein-level false-discovery rates (FDR). Processed data was searched against a database composed of computationally predicted open reading frames (ORFs) of the metagenomes retrieved from the same culture.

**GC-MS analysis of low molecular weight aromatic compounds**

Basic experimental procedures were carried out following the methodology of Raj et al.[1] with some modifications. Specifically, aliquots (2 mL) of enrichment cultures or controls were sampled at 0 d and 30 d and spiked with a surrogate standard (ethyl vanillin). After acidification with HCl (pH<2) and oversaturation with NaCl, low-molecular-weight (LMW) aromatic compounds were extracted from the aqueous phase with 1 mL ethyl acetate three times. The extracts were pooled, dewatered over anhydrous $Na_2SO_4$, and then concentrated under nitrogen. Before GC-MS analysis,

LMW compounds were converted to trimethylsilyl derivatives with N,O-bis-(trimethylsilyl) trifluoroacetamide (BSTFA) and pyridine (70°C, 45 min).

Silylated compounds were identified and quantified on a Trace 1310 gas chromatograph coupled to a TSQ8000 mass spectrometer (Thermo Fisher Scientific, USA) using a HP-5MS capillary column (30 m × 0.25 mm i.d., 0.25 μm film thickness). The temperature programming was as follows: 2 min at 65°C, 6°C min$^{-1}$ to 300°C, and 20 min at 300°C. The injector temperature was set at 300°C. The transfer line and ion source temperatures were maintained at 300 and 290°C respectively. Helium was used as carrier gas at a flow rate of 1 mL min$^{-1}$. The mass spectrometer was operated in electron impact (EI) mode at 70 eV. EI mass spectra were recorded from 50 to 650 m/z in full-scan mode. Identification of compounds was achieved by comparing the mass spectra with that of the NIST library and by comparing the retention time with that of available authentic compounds. Key compounds (vanillin, acetovanillone, vanillic acid, homovanillic acid, syringic acid, guaiacol, protocatechoic acid, and catechol) were quantified by comparing with surrogate standards (ethyl vanillin) to account for compound loss during extraction procedures. External quantification standards were used to normalize the response factor for different compounds separately.

**Heterologous protein production of MtgC and MtgB_2:**

The gene encoding the methyltransferase I (MtgB_2) was amplified from the DL1YTT001 culture with primers F1/R1 (F1: 5’-GGTACCCTCGAGGGATCCATGAAGTTTGGAATGTTCATTTATG-3’; R1: 5’-CAGGTCGACAAGCTTTTAGTCTAGTATACTTTCCCACTTGTC-3’) for cloning in expression vector pCold-TF inserting an N-terminal Strep tag via the reverse primer. For cloning of the MtgB_2 genes into pCold-TF, primers included BamHI and HindIII restriction sites to insert the purified PCR products into the plasmid.

For Co(I) production, the activating enzyme (AE) gene of Acetobacterium dehalogenans DSM 11527 (GenBank accession no. ACJ01666.1) was synthesized. The AE gene and corrinoid protein (MtgC) gene was amplified with primers AE-F/R (AE-F: 5’-gCCgCgCggCAGCCATATGATGTCATCTTTGAATACT-3’ AE-R: 5’-CGAGTGCGGCCGCAAGCTTTTATTTCATTTCATTTTG-3’) and F2/R2 (F2: 5’-

GCCGCGCGGCAGCCATATGATGTCTTGGTTAAAATCTATGATG-3’; R2: 5’-CGAGTGCGGCCGCAAGCTTTTATTTGGATGCCATTGCTTTTTT-3’) for cloning in expression vector pET-30a inserting an N-terminal Strep tag via the reverse primer. For cloning of AE and MtgC genes into pET-30a, primers included NdeI and and HindIII restriction sites to insert the purified PCR products into the plasmid.

PCR was performed with PrimeSTAR® Max DNA Polymerase (Takara) according to manufacturer’s instructions. PCR products were purified using an Agarose Gel DNA Fragment Recovery Kit Ver.2.0 (TaKaRa, Dalian, China). The purified PCR products were ligated into expression vectors by using ligase Exnase® II (Vazyme Biotech Co., Ltd, Nanjing, China). *E. coli* DH5α (Takara, Dalian, China) was used for plasmid transformation. Plasmid DNA for cloning and sequencing was all prepared using the Plasmid Mini Kit I (Omega).

For production of MtgB_2, MtgC and AE, the plasmids were transformed into *E. coli* BL21(DE3). Freshly transformed *E.coli* BL21 was grown aerobically in a 50ml overnight culture (Luria-Bertani medium 100 µg/ml ampicillin, 37°C). This culture was transferred into 1L of the same medium, which was induced with 0.5 mM IPTG. After growth for an additional 18 h, cells were harvested by centrifugation at 8000 rpm/min for 10 min at 4 °C. All following steps were performed in an anoxic environment in an anaerobic chamber with anoxic buffers and solutions. All solutions were made anoxic by sparging for 10 min with nitrogen gas. Pelleted cells harvested above were resuspended in 40 mL lysis buffer (20 mmol/L Tris-HCl, pH 7.6, 150 mmol/L NaCl, 5% glycerol) and lysed by sonication (4 s pulse, 4 s pause, 600W; 20 min). After removal of insoluble cell materials by centrifugation (10000 rpm/min for 30 min at 4 °C), the supernatants were purified by immobilized $Ni^{2+}$ affinity chromatography. After loading the supernatants onto the column pre-equilibrated with lysis buffer, the resin was washed with lysis buffer containing 20 mM imidazole. The target proteins were then eluted with lysis buffer containing 200 mM imidazole. Afterwards, the buffer was exchanged with 20 mmol/L Tris-HCl (pH 7.6), 1 mmol/L DTT, 150 mmol/L NaCl and 5% glycerol using 10 kDa ultrafiltration units (Amicon Ultra-15 filter) and stored in small aliquots at -20°C in an anaerobic bottle ($N_2/H_2$ (95%/5%)). The purity of eluted fractions was confirmed by 15% SDS-PAGE.

The protocol for reconstitution of MtgC with cobalamin was adapted from Schilhabel et al.[2] and Kurth et al.[3]. 1.5 ml (30 mg) anaerobic protein solution was added to 8.5 ml refolding solution containing 1 mM DTT and incubated in the dark at 4°C for 16 h in a glass bottle closed with a rubber stopper. The refolding solution contained 50 mM Tris HCl, 3.5 M betaine and 1 mM hydroxocobalamin, and pH was adjusted to 7.6. The protein solution then was incubated for 16 h at 4 °C in the dark. Afterwards, the buffer was exchanged several times with Tris HCl (pH 7.5) and 1 mM DTT using 5 kDa ultrafiltrationn units (Amicon Ultra-15 Centrifugal Filter Units, Merck) until the cobalt-containing permeate appeared visibly clear instead of red. Protein was stored anaerobically in 2 ml glass vials sealed with air-tight rubber stoppers.

Table S1 The relative abundance of microbial populations based on 16S rRNA gene-tag sequencing analysis before and after antibiotic treatment.

| taxonomy | before antibiotics | after antibiotics |
|---|---|---|
| DL1YTT001 (*Bathyarchaeia*) | 60.10% | 88.89% |
| *Desulfobacteraceae* (*Desulfobacterota*) | 13.22% | 0.63% |
| *Spirochaetaceae* (*Spirochaetota*) | 10.50% | 1.68% |
| *Desulfovibrionaceae* (*Desulfobacterota*) | 5.46% | 0.95% |
| *Deferribacteraceae* (*Deferribacterota*) | 2.18% | 0.06% |
| *Synergistaceae* (*Synergistota*) | 2.05% | 0.14% |
| *Christensenellaceae* (*Firmicutes*) | 1.64% | 0.45% |
| *Syntrophotaleaceae* (*Desulfobacterota*) | 0.82% | 1.84% |
| *Methanocorpusculaceae* (*Euryarchaeota*) | 0.67% | 0.04% |
| LCP-89 | 0.65% | 0.17% |
| *Desulfatiglandaceae* (*Desulfobacterota*) | 0.54% | 0.00% |
| *Alkalibacteraceae* (*Firmicutes*) | 0.06% | 2.54% |
| Other | 1.28% | 2.61% |

Table S2 Overview of the MAG of strain DL1YTT001.

| | DL1YTT001 |
|---|---|
| Completeness, % | 99.22 |
| Contamination, % | 2.8 |
| Genome size, Mb | 1.81 |
| Number of contigs | 43 |
| N50 value, bp | 99919 |
| Mean contig length, bp | 42151 |
| GC content, % | 40 |
| Predicted genes | 1905 |

Table S3 Transcription and translation of central carbon metabolism genes. +, detected; -, undetected.

Table S4 Distribution of methyltransferase genes in Bathyarchaeial MAGs. C indicate MAG contain the methyltransferase gene cluster; G indicate MAG contain the gene of methyltransferase 1. NA indicate no methyltransferase gene or gene cluster was found in MAG.

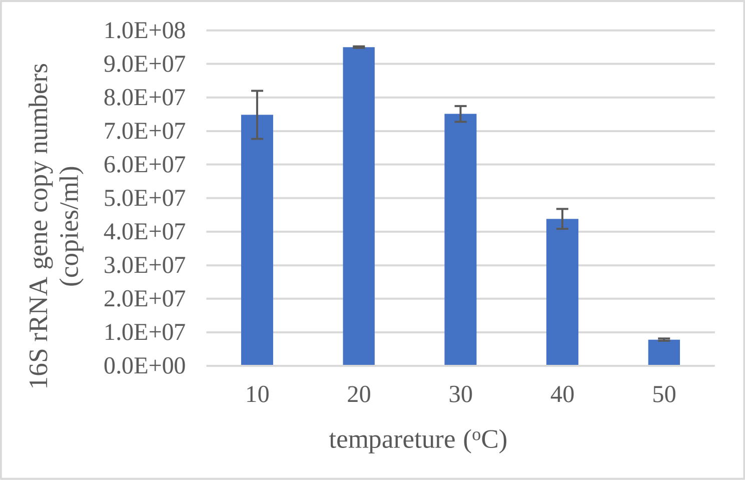


Fig. S1 The effect of incubation temperature on growth of strain DL1YTT001. Error bars indicate standard deviations of duplicate determinations.

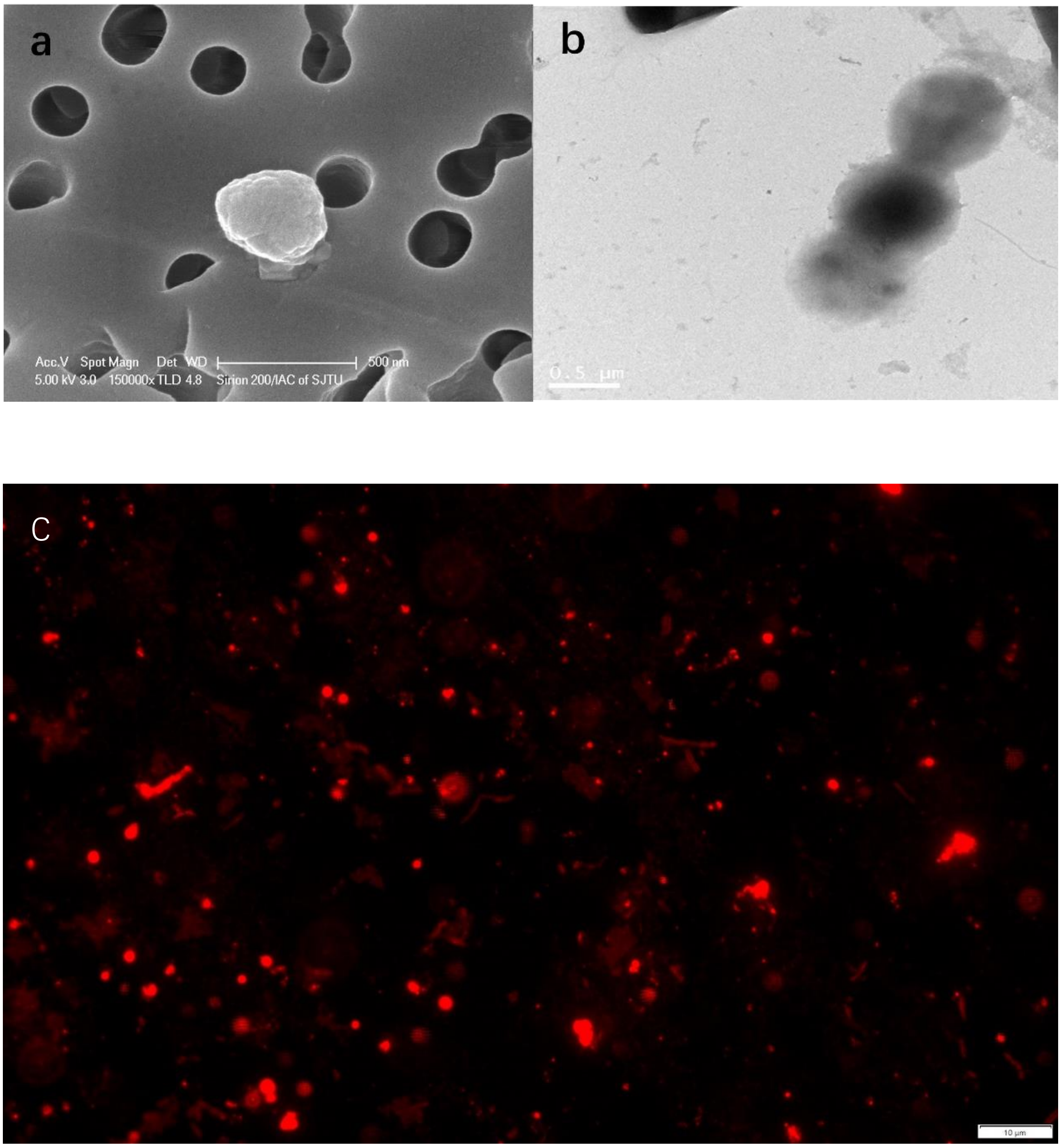
a
Acc.V Spot Magn Det WD 500 nm
5.00 kV 3.0 150000x TLD 4.8 Sirion 200/IAC of SJTU
b
0.5 μm
C
10 μm

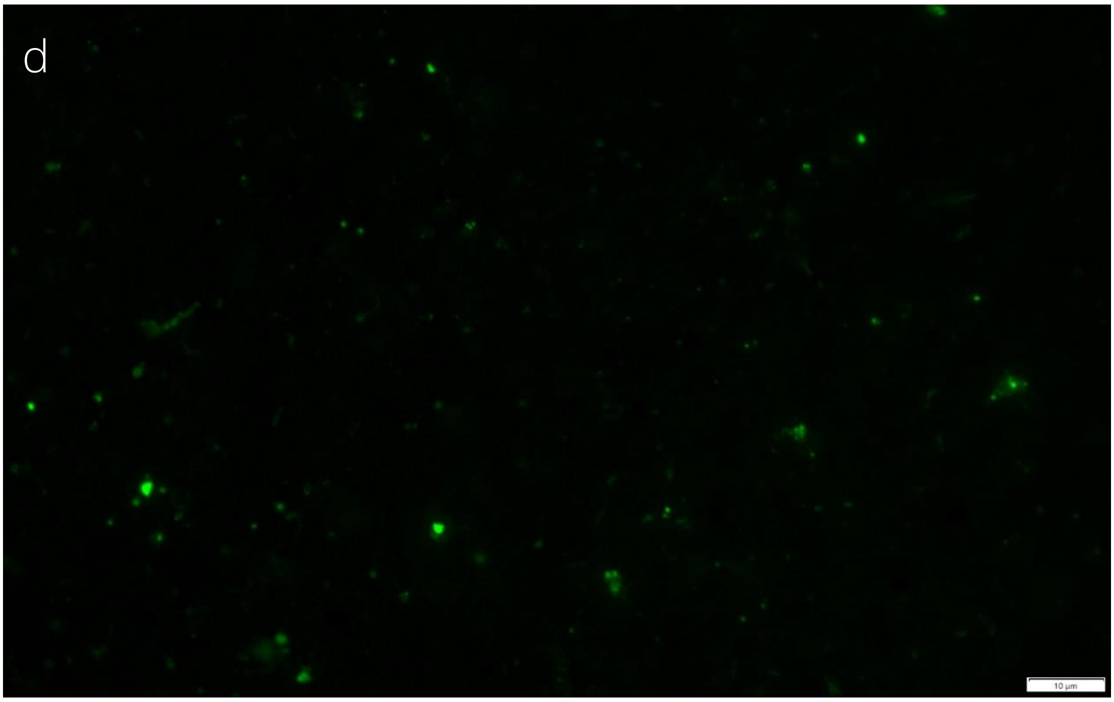
d
10 μm

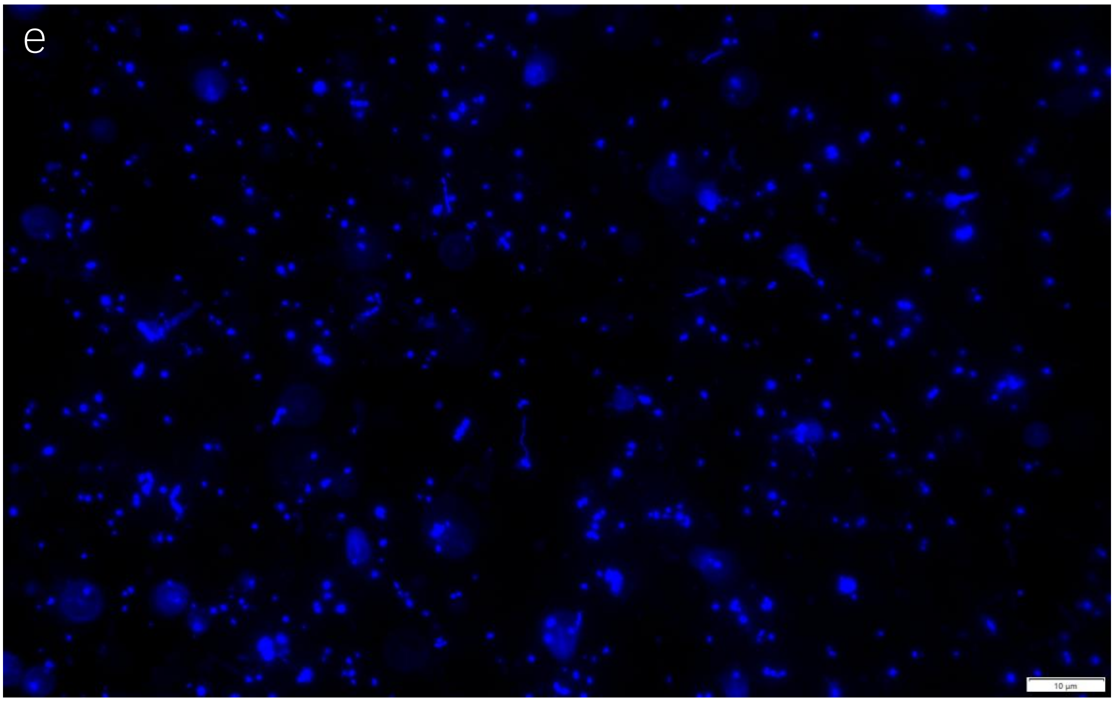
e
10 μm

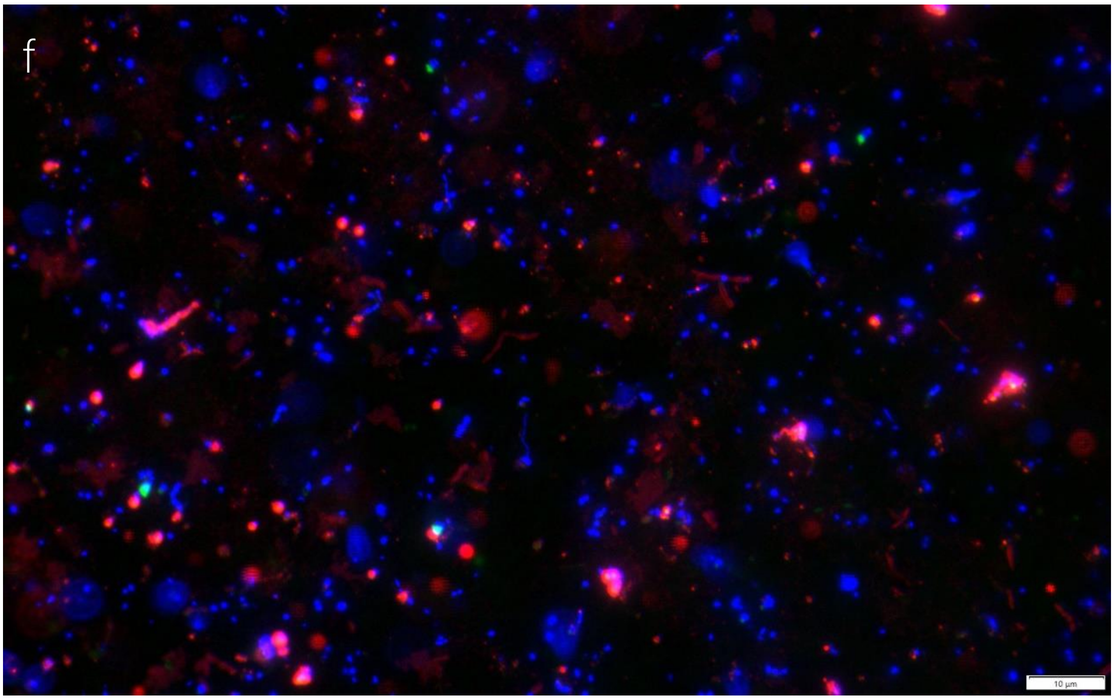


Fig S2 Scanning Electron Microscope (SEM) (a), Transmission Electron Microscope (TEM) (b), and Fluorescence in situ hybridization (FISH) images of strain DL1YTT001 (c-f). For FISH, DL1YTT001 cells were hybridized with a *Bathyarchaeia* 16S rRNA-targeted probe (labeled by Alexa Fluor 594, red fluorescence) (c), and bacterial cells were hybridized with a Bacterial 16S rRNA-targeted probe (labeled by Alexa Fluor 488, green fluorescence) (d). After hybridization, the cells were counterstained with DAPI (blue fluorescence) (e). A composite image of all three filters (f).

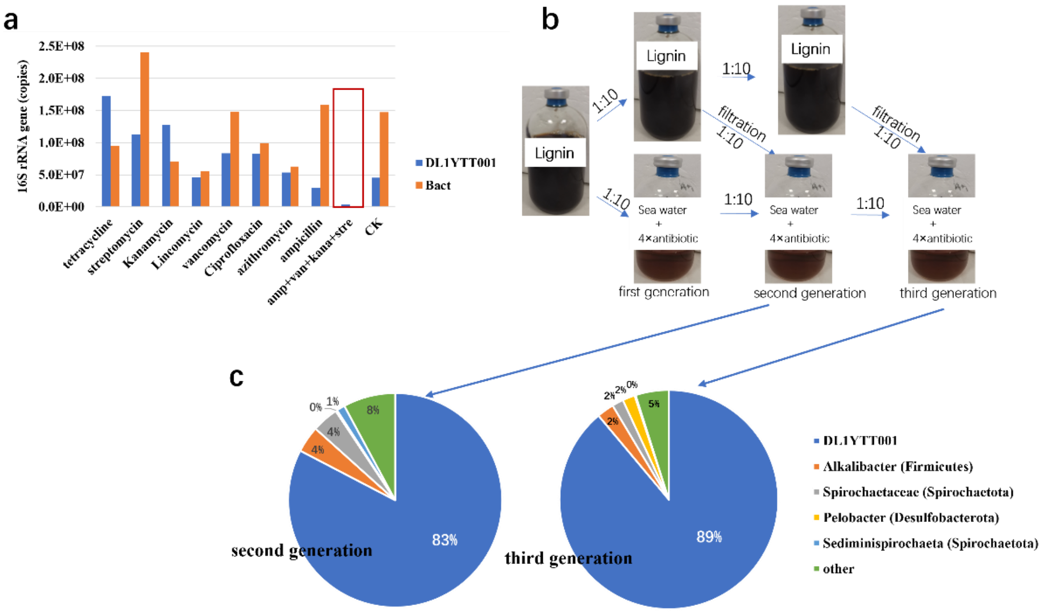


Fig. S3 Schematic diagram of the purification process for strain DL1YTT001. (a) Inhibition effects of individual antibiotics (tetracycline, streptomycin, Kanamycin, Lincomycin, vancomycin, Ciprofloxacin, azithromycin, ampicillin), and an antibiotic cocktail (ampicillin, vancomycin, kanamycin, and streptomycin) on DL1YTT001 and Bacterial growth. Each antibiotic was supplemented at a final concentration of 50 µg/ml. (b) The antibiotic enrichment process for DL1YTT001: The DL1YTT001 culture supplemented with antibiotic cocktail was transferred to media containing supernatant from the same culture (taken prior to antibiotic treatment) in order to supplement putative bacterial metabolites in the supernatant. The antibiotic cocktail was also supplemented to inhibit Bacterial growth. (c) The relative abundance of microbial populations in the second and third generations based on 16S rRNA gene-tag sequencing analysis.

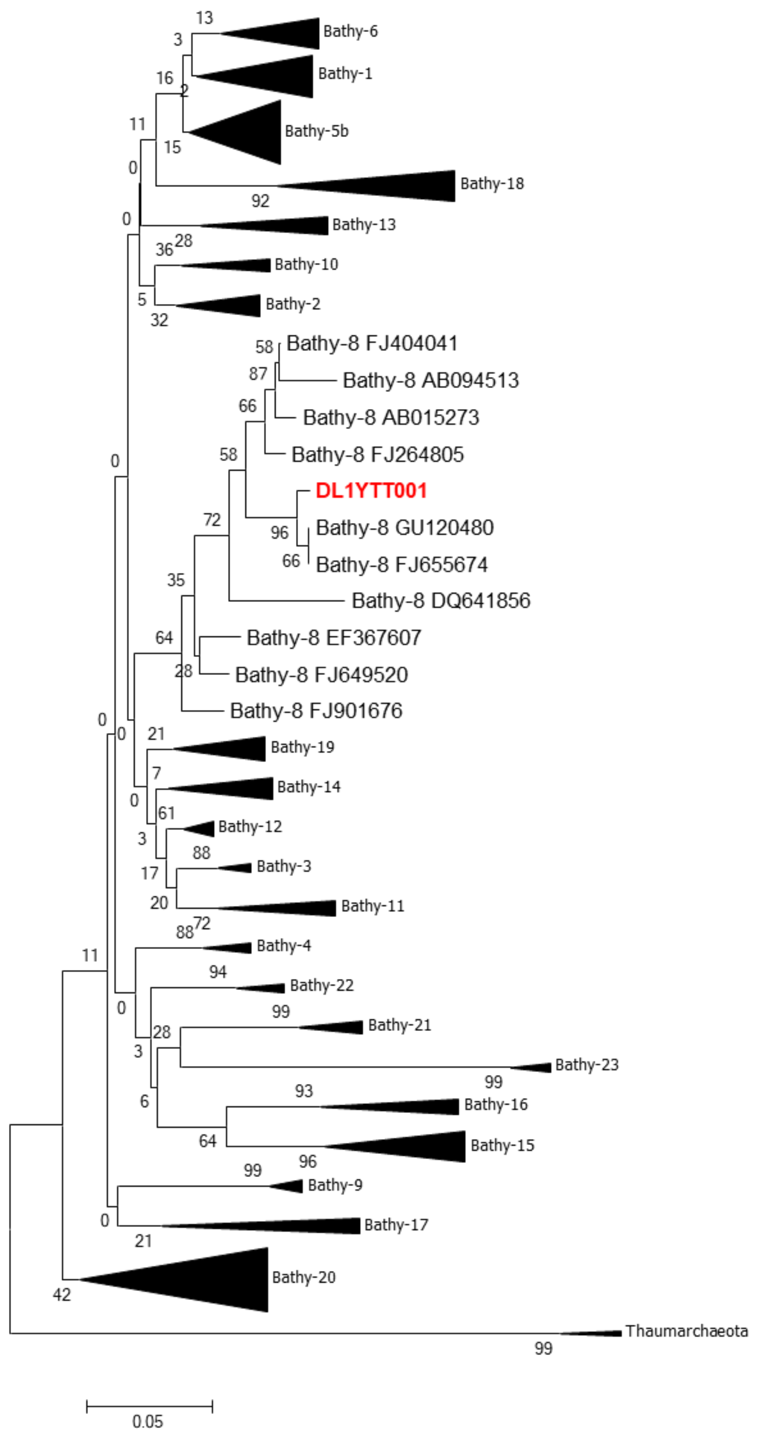


Fig. S4 Maximum-likelihood phylogeny of *Bathyarchaeia* 16S rRNA genes. Bootstrap values were calculated from 1,000 iterations using Mega (MEGA-X).

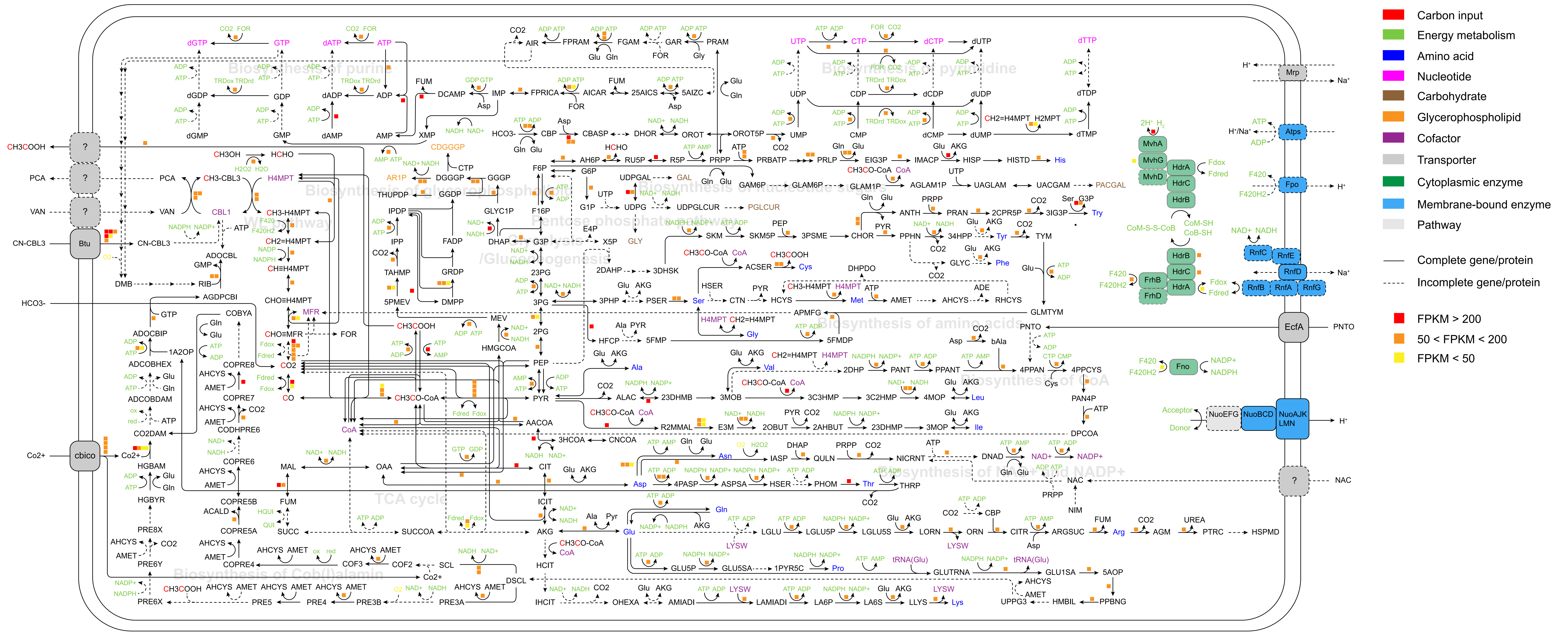

Carbon input
Energy metabolism
Amino acid
Nucleotide
Carbohydrate
Glycerophospholipid
Cofactor
Transporter
Cytoplasmic enzyme
Membrane-bound enzyme
Pathway
Complete gene/protein
Incomplete gene/protein
FPKM > 200
50 < FPKM < 200
FPKM < 50
Biosynthesis of purine
Biosynthesis of pyrimidine
Biosynthesis of glycerophospholipid
Biosynthesis of nucleotide sugars
Pentose phosphate pathway
Glycolysis /Gluconeogenesis
WL pathway
Biosynthesis of amino acids
Biosynthesis of CoA
Biosynthesis of NAD+ and NADP+
TCA cycle
Biosynthesis of Cob(I)alamin
Btu
cbico
EcfA
Mrp
Atps
Fpo
RnfC
RnfE
RnfD
RnfB
RnfA
RnfG
MvhA
MvhG
MvhD
HdrA
HdrC
HdrB
FrhB
FrhD
Fno
NuoEFG
NuoBCD
NuoAJK LMN
CH3COOH
PCA
VAN
CN-CBL3
HCO3-
Co2+
PNTO
NAC
H⁺
Na⁺

Fig. S5 Overview of the metabolic network of strain DL1YTT001. It includes the assimilation pathway of methyl groups derived from methoxylated aromatic compounds, glycolysis/gluconeogenesis, the pentose phosphate pathway, the Wood-Ljungdahl pathway, the TCA cycle, biosynthesis pathways of biomass precursors (amino acids, nucleic acids, glycerophospholipids, and polysaccharides), cofactors (Cob(I)alamin, $NAD^+$/NADP+, Coenzyme A, and methanofuran), and several key membrane transporters. Solid lines and dotted lines refer to genes present and missing in the DL1YTT001 genome, respectively. Colored squares refer to levels of gene transcription.

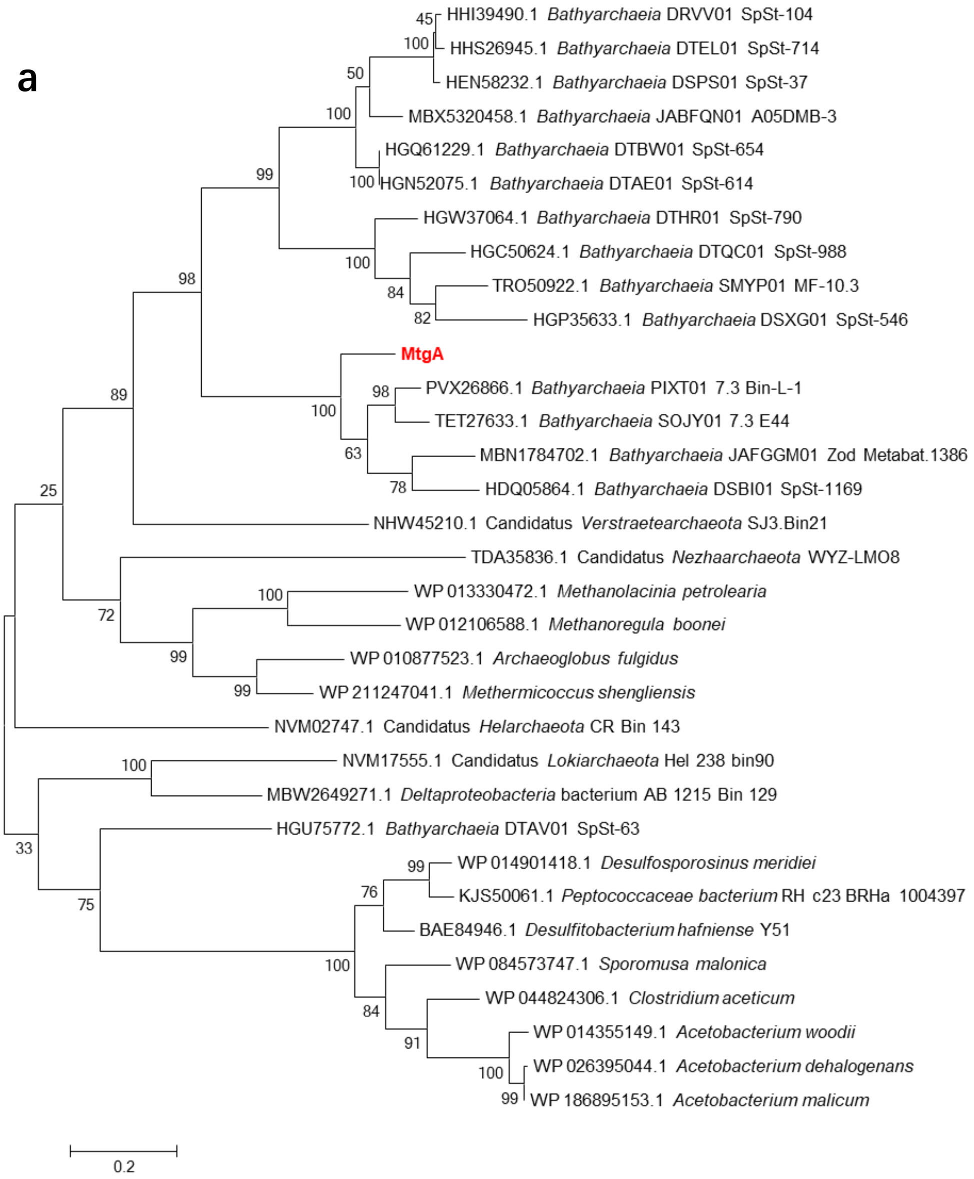
a
HHI39490.1 Bathyarchaeia DRVV01 SpSt-104
HHS26945.1 Bathyarchaeia DTEL01 SpSt-714
HEN58232.1 Bathyarchaeia DSPS01 SpSt-37
MBX5320458.1 Bathyarchaeia JABFQN01 A05DMB-3
HGQ61229.1 Bathyarchaeia DTBW01 SpSt-654
HGN52075.1 Bathyarchaeia DTAE01 SpSt-614
HGW37064.1 Bathyarchaeia DTHR01 SpSt-790
HGC50624.1 Bathyarchaeia DTQC01 SpSt-988
TRO50922.1 Bathyarchaeia SMYP01 MF-10.3
HGP35633.1 Bathyarchaeia DSXG01 SpSt-546
MtgA
PVX26866.1 Bathyarchaeia PIXT01 7.3 Bin-L-1
TET27633.1 Bathyarchaeia SOJY01 7.3 E44
MBN1784702.1 Bathyarchaeia JAFGGM01 Zod Metabat.1386
HDQ05864.1 Bathyarchaeia DSBI01 SpSt-1169
NHW45210.1 Candidatus Verstraetearchaeota SJ3.Bin21
TDA35836.1 Candidatus Nezhaarchaeota WYZ-LMO8
WP 013330472.1 Methanolacinia petrolearia
WP 012106588.1 Methanoregula boonei
WP 010877523.1 Archaeoglobus fulgidus
WP 211247041.1 Methermicoccus shengliensis
NVM02747.1 Candidatus Helarchaeota CR Bin 143
NVM17555.1 Candidatus Lokiarchaeota Hel 238 bin90
MBW2649271.1 Deltaproteobacteria bacterium AB 1215 Bin 129
HGU75772.1 Bathyarchaeia DTAV01 SpSt-63
WP 014901418.1 Desulfosporosinus meridiei
KJS50061.1 Peptococcaceae bacterium RH c23 BRHa 1004397
BAE84946.1 Desulfitobacterium hafniense Y51
WP 084573747.1 Sporomusa malonica
WP 044824306.1 Clostridium aceticum
WP 014355149.1 Acetobacterium woodii
WP 026395044.1 Acetobacterium dehalogenans
WP 186895153.1 Acetobacterium malicum
0.2

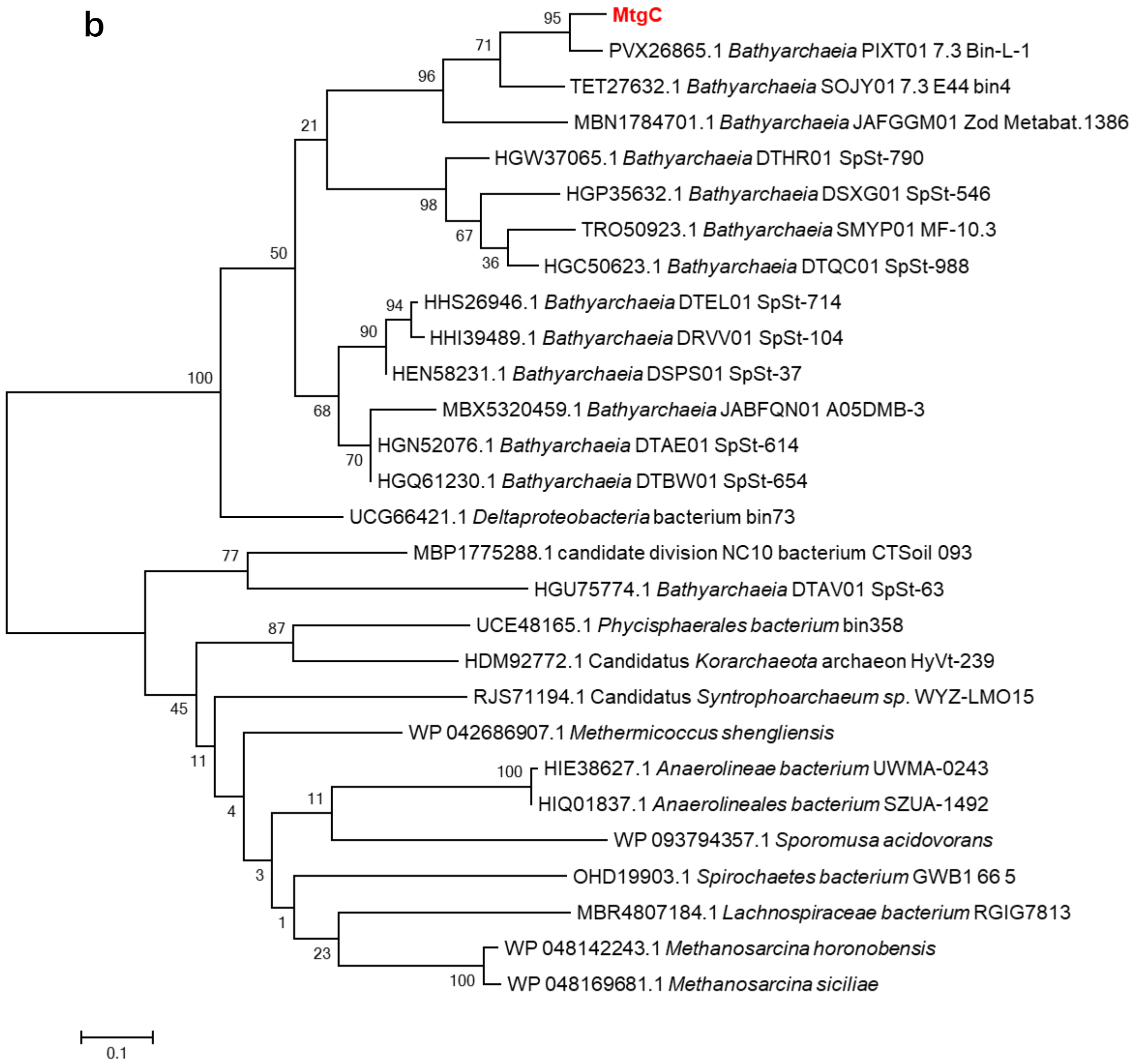


Fig. S6 Maximum-likelihood phylogeny of MtgA (methyltransferase 2, MT2) (a), and MtgC (corrinoid protein, CP) (b) within the class Bathyarchaeia. Bootstrap values were calculated from 1,000 iterations using Mega (MEGA-X).

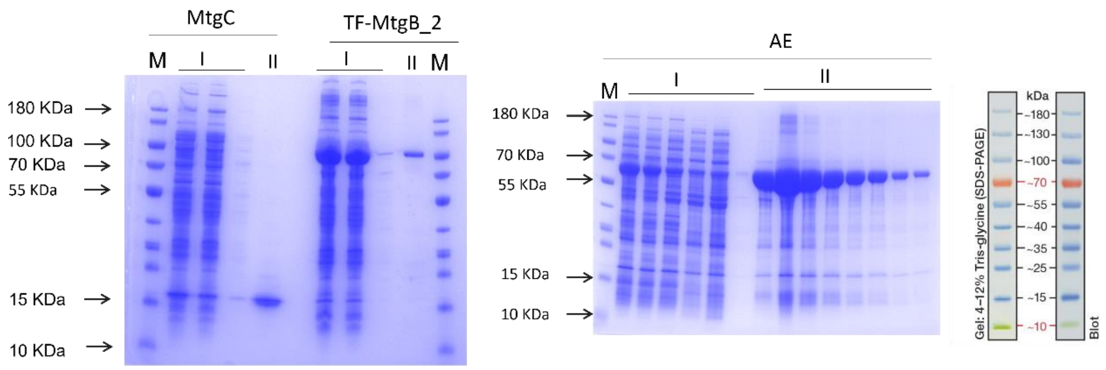


Fig. S7 SDS-PAGE of heterologously expressed methyltransferase 1 (MtgB_2), corrinoid protein (MtgC), and activating enzyme (AE). MtgB_2 and MtgC were from DL1YTT001, and AE was from *Acetobacterium dehalogenans* DSM 11527 (GenBank accession no. ACJ01666.1). M, molecular mass marker; I, crude enzyme solution; II, the purified protein samples.

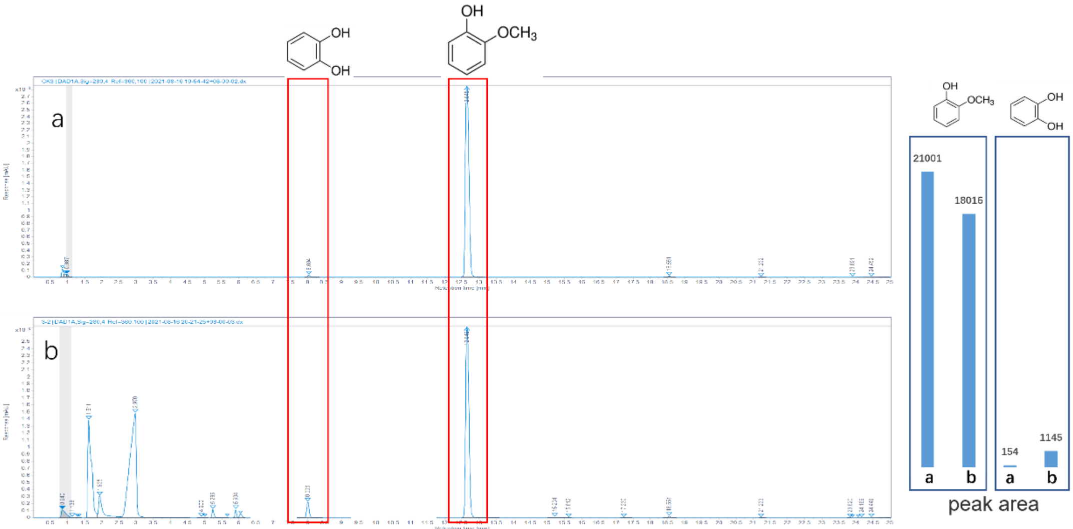


Fig. S8 HPLC analysis of the bioconversion of methoxylated aromatic compounds catalyzed by MtgB_2. The MtgB_2 activity was determined in enzyme activity assays using 2.3 mM guaiacol as a substrate. 50 µl of the sample were collected before (a) and after (b) addition of the MtgB_2 for analysis of methoxy aromatic compounds and products by HPLC. The retention time of standards of guaiacol and catechol were determined to be 12.65 min and 8.00 min, respectively. After addition of the MtgB_2 (b), the guaiacol peak decreased by around 14.01% along with an increase of the catechol peak.

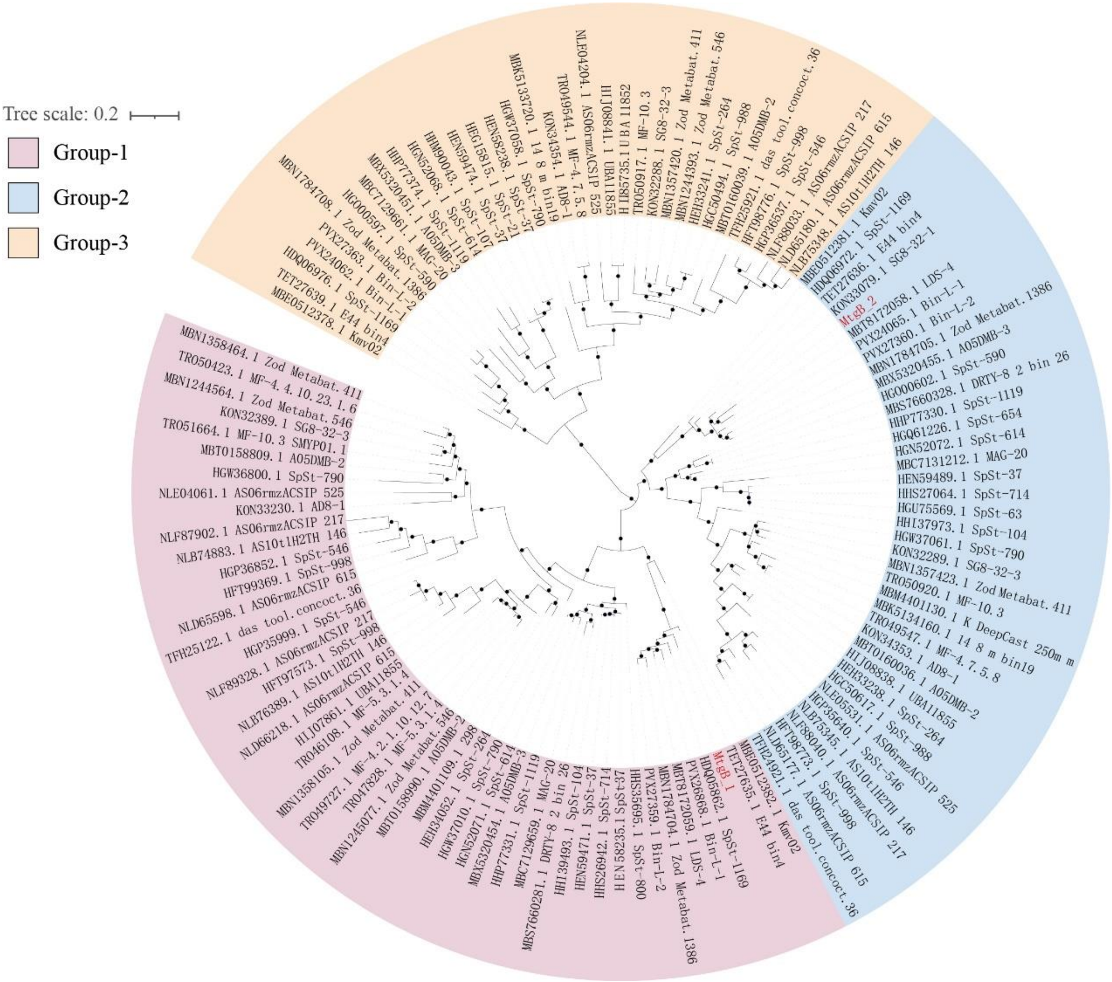


Fig. S9 Maximum-likelihood phylogeny of MtgB (methyltransferase 1, MT1) within the class *Bathyarchaeia*. Bootstrap values were calculated from 1,000 iterations using Mega (MEGA-X). Bootstrap values of >70% are labelled with black dots.
